# Phosphoinositide Depletion and Compensatory β-adrenergic Signaling in Angiotensin II-Induced Heart Disease: Protection Through PTEN Inhibition

**DOI:** 10.1101/2025.02.23.639781

**Authors:** Taylor L. Voelker, Maartje Westhoff, Silvia G. del Villar, Phung N. Thai, Nipavan Chiamvimonvat, Madeline Nieves-Cintrón, Eamonn J. Dickson, Rose E. Dixon

## Abstract

Contractile dysfunction, hypertrophy, and cell death during heart failure are linked to altered Ca^2+^ handling, and elevated levels of the hormone angiotensin II (AngII), which signals through G_q_-coupled AT_1_ receptors, initiating hydrolysis of PIP_2_. Chronic elevation of AngII contributes to cardiac pathology, but the mechanisms linking sustained AngII signaling to heart dysfunction remain incompletely understood. Here, we demonstrate that chronic AngII exposure profoundly disrupts cardiac phosphoinositide homeostasis, triggering a cascade of cellular adaptations that ultimately impair cardiac function. Using *in vivo* AngII infusion combined with phospholipid mass spectrometry, super-resolution microscopy, and functional analyses, we show that sustained AngII signaling reduces PI(4,5)P_2_ levels and triggers extensive redistribution of Ca_V_1.2 channels from t-tubules to various endosomal compartments. Despite this t-tubular channel loss, enhanced sympathetic drive maintains calcium currents and transients through increased channel phosphorylation via PKA and CaMKII pathways. However, this compensation proves insufficient as cardiac function progressively declines, marked by pathological hypertrophy, t-tubule disruption, and diastolic dysfunction. Notably, we identify depletion of PI(3,4,5)P_3_ as a critical mediator of AngII-induced cardiac pathology. While preservation of PI(3,4,5)P_3_ levels through PTEN inhibition did not prevent cellular remodeling or calcium handling changes, it protected against cardiac dysfunction, suggesting effects primarily through reduction of fibrosis. These findings reveal a complex interplay between phosphoinositide signaling, ion channel trafficking, and sympathetic activation in AngII-induced cardiac pathology. Moreover, they establish maintenance of PI(3,4,5)P_3_ as a promising therapeutic strategy for hypertensive heart disease and as a potential protective adjunct therapy during clinical AngII administration.

Graphical Abstract.
Graphical summary of chronic angiotensin II (AngII) signaling effects on cardiac calcium handling and fibrosis.
Chronic AngII signaling through AT1R activates G_q_-coupled signaling, leading to PI(4,5)P_2_ depletion that destabilizes Ca_V_1.2 in the plasma membrane (PM), triggering their endocytosis and reduced channel numbers at the PM. The remaining Ca_V_1.2 channels and RyR2 undergo compensatory phosphorylation by CaMKII and PKA, triggered by sympathetic activation (β-AR signaling), leading to enhanced calcium-induced calcium release (CICR). Meanwhile, AngII promotes fibroblast-to-myofibroblast transition via M1 macrophage phenotype activation, increasing cardiac fibrosis. PTEN inhibition preserves PIP_3_ levels and promotes anti-inflammatory M2 macrophage activation, resulting in reduced fibrosis. These findings reveal a complex interplay between cardiac phosphoinositide signaling, calcium handling, and fibrotic remodeling with chronic AngII. AC, adenylyl cyclase; β-AR, beta-adrenergic receptor; DAG, diacylglycerol; IP_3_, inositol trisphosphate; PLC, phospholipase C; PKA, protein kinase A; PTEN, phosphatase and tensin homolog.

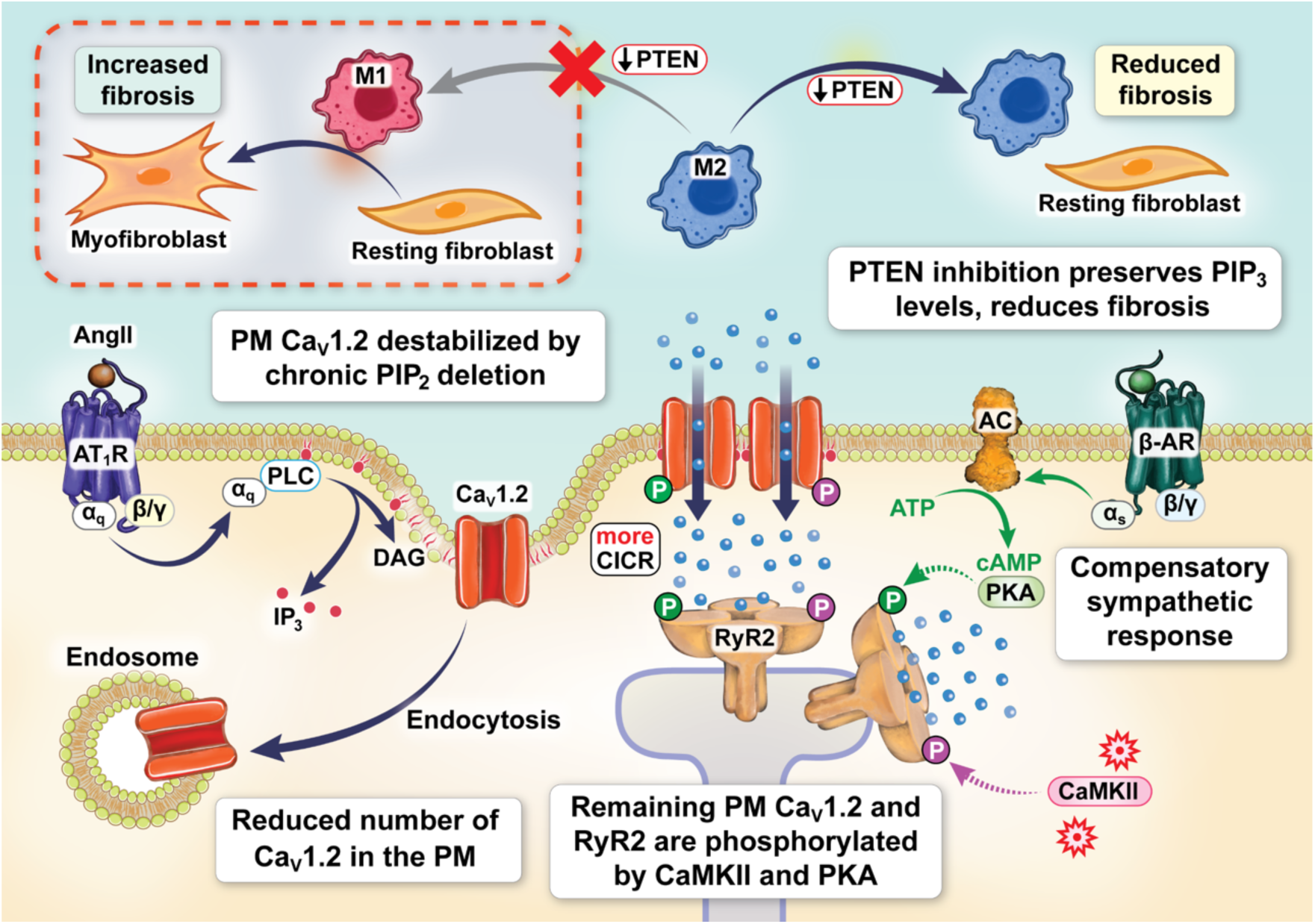

## Introduction

Angiotensin II (AngII), a key neurohormone of the Renin-Angiotensin-Aldosterone-System (RAAS), functions physiologically as a potent vasoconstrictor in blood pressure regulation while contributing pathologically to hypertension^1, 2^ and heart failure (HF)^3–5^. In HF, sustained elevation of AngII contributes to pathological cardiac remodeling through promotion of fibrosis, contractile dysfunction, hypertrophy, and cell death^6–11^. HF is also associated with altered Ca^2+^ handling, and dysregulation of Ca_V_1.2 channel function, trafficking, and localization^2, 12–14^. Despite this correlative association, there is no clear consensus on the functional effects of chronic elevations in AngII on cardiac Ca_V_1.2 expression, trafficking, and activity.

AngII signals through both G_q_-coupled AngII type 1 receptors (AT_1_R), and G_i_-coupled AngII type 2 receptors (AT_2_R) in the heart^15^. Notably mice with cardiac-specific inactivation of G_q_^16^ or deletion of phospholipase Cε (PLCε, an enzyme downstream from G_q_) fail to develop hypertrophy in response to pressure overload^17^. These studies suggest the AngII/AT_1_R/G_q_/PLCε signaling axis plays a causative role in cardiac hypertrophy. Strengthening that idea, cardiac-specific AT_1_R overexpression leads to spontaneous hypertrophy and HF in the absence of hypertension, suggesting that although hypertension can form a substrate for HF, enhanced cardiac AngII/AT_1_R signaling is sufficient to drive cardiac remodeling and precipitate this pathology^6, 18–21^. Conversely, AngII infusion-induced cardiac hypertrophy is similar in WT and AT_2_R overexpressing mice^22, 23^, and the hypertrophic response to aortic-banding can be reduced by AT_1_R antagonists but is unaltered in AT_2_R knockouts^24^. Thus, AT_2_R signaling is not thought to alter cardiac hypertrophy.

Graded cardiac Ca_V_1.2 channel deletion leads to spontaneous hypertrophy and HF in mice^25^, so alterations to trafficking or expression of Ca_V_1.2 itself could also act as a substrate for HF. The number and activity of Ca_V_1.2 channels in the sarcolemma tunes *I*_Ca_, Ca^2+^-induced Ca^2+^ release, and inotropy^26–28^. We recently reported that *acute* AngII applied to isolated adult mouse ventricular myocytes stimulated measurable hydrolysis of the plasma membrane phospholipid phosphatidylinositol (4,5)-bisphosphate (PI(4,5)P_2_) via the AngII/AT_1_R/G_q_/PLCε signaling axis^29^. This reduction in PI(4,5)P_2_ (henceforth referred to as PIP_2_) was shown to trigger dynamin-mediated endocytosis of Ca_V_1.2 channels from the t-tubule sarcolemma of cardiomyocytes, resulting in reduced whole-cell Ca^2+^ currents (*I*_Ca_) and diminished Ca^2+^ transient amplitudes. Those findings provide an explanation as to why voltage-gated Ca^2+^ channel currents decrease upon activation of G_q_-coupled pathways and are preserved with PIP_2_ supplementation, as others have reported^30–32^.

In this study, we investigated how sustained AngII signaling—a hallmark of hypertension and HF—alters the cardiac phospholipid landscape and its subsequent effects on Ca_V_1.2 channel trafficking, expression, and function, as well as EC-coupling and overall cardiac performance. Using *in vivo* osmotic minipump delivery, we hypothesized that chronic AngII signaling would cause prolonged PIP_2_ and Ca_V_1.2 deficits, leading to reduced cardiac function. Through phospholipid mass spectrometry, electrophysiology, functional analyses, and advanced imaging approaches, we discovered that chronic AngII infusion disrupted PIP_2_ and multiple phosphoinositide species. This profound lipid remodeling was accompanied by cytoarchitectural changes and reduced t-tubular Ca_V_1.2 expression and clustering. While enhanced sympathetic drive functionally upregulated the depleted Ca_V_1.2 population to maintain Ca^2+^ signaling, echocardiography revealed persistent cardiac dysfunction in AngII-infused mice. Notably, among the altered phosphoinositides, we identified PIP_3_ depletion as a critical mediator of AngII pathology. Pharmacological inhibition of PTEN, a 3’-lipid phosphatase that dephosphorylates PI(3,4,5)P_3_ to form PI(4,5)P_2_, prevented AngII-induced cardiac dysfunction, suggesting maintenance of phosphoinositide homeostasis as a potential therapeutic strategy.

## Methods

### Osmotic Mini-Pump Infusion

Male C57Bl/6J mice (8-10 weeks old, The Jackson Laboratory, Sacramento, CA, USA) underwent surgery to implant subcutaneous osmotic minipumps (Alzet, Durect Corp., Cupertino, CA) delivering either Ang II (MilliporeSigma, Rockville, MD, USA; 1 mg/kg/day), BpV(phen) (MilliporeSigma; 17 µg/kg/day), a combination of both, or sterile saline (Covidien, Dublin, Ireland). After 1 week, mice were euthanized by intraperitoneal injection of pentobarbital solution (Beuthanasia-D Special; Merck Animal Health, Madison, NJ, USA), and hearts were collected for subsequent experiments and analysis.

### Isolation of mouse ventricular myocytes

Upon removal from the thorax, hearts were plunged into ice-cold digestion buffer (130 mM NaCl, 5 mM KCl, 3 mM Na-pyruvate, 25 mM HEPES, 0.5 mM MgCl_2_, 0.33 mM NaH_2_PO_4_, and 22 mM glucose) supplemented with 150 μM EGTA. Individual myocytes were isolated via retrograde Langendorff perfusion at 37 °C. Following blood clearance, hearts were perfused with digestion buffer containing 50 μM CaCl_2_ (Thermo Fisher Scientific, Rockford, IL, USA), 0.04 mg/ml protease (MilliporeSigma, Rockville, MD, USA), and 1.4 mg/ml type 2 collagenase (Worthington Biochemical, Lakewood, NJ, USA) until pale and soft to the touch. Ventricles were then minced and further digested in buffer containing collagenase (0.96 mg/ml), protease (0.04 mg/ml), CaCl_2_ (100 μM), and BSA (10 mg/ml). Individual myocytes were dispersed by gentle trituration.

### Single-Molecule Localization Microscopy (SMLM)

No. 1.5 coverslips were sonicated in 2 N NaOH (20 min), rinsed with deionized water, and coated with poly-L-lysine (0.01%; MilliporeSigma) and laminin (20 μg/ml; Life Technologies, Carlsbad, CA, USA). Isolated myocytes were plated and allowed to adhere (37 °C, 45 min) before fixation in ice-cold methanol (Thermo Fisher Scientific, −20 °C, 5 min). After PBS washes, samples were blocked (20% SEA Block, 0.5% Triton X-100 in PBS, 1 hr) and incubated with anti-Ca_V_1.2 antibody (CACNA1C, ACC-003, Alomone Labs, Jerusalem, Israel, 1:300, overnight at 4 °C). Following PBS washes, samples were labeled with Alexa Fluor 647-conjugated secondary antibody (Life Technologies, 1:1000, 1hr at RT). Coverslips were mounted onto depression glass slides in oxygen-scavenging buffer (50 mM Tris pH 8.0, 10 mM NaCl, 0.56 mg/ml glucose oxidase, 34 μg/ml catalase, 10% glucose, 100 mM MEA). To exclude oxygen, the edges of the coverslips where sealed using Twinsil dental glue (Picodent, Wipperfürth, Germany) and aluminum tape (T205-1.0 - AT205; Thorlabs Inc., Newton, NJ, USA). Images were acquired using either a Leica 3D-GSD-SR or DMi8 microscope (Leica Microsystems, Wetzlar, Germany) in TIRF mode with 150 nm penetration depth using 160×/1.43 NA objectives. For the 3D-GSD-SR, a 642 nm/500 mW laser and iXon3 EM-CCD camera (Andor, Oxford Instruments, Oxfordshire, UK) were used for excitation and photon detection; for the DMi8, a 638 nm/150 mW laser and ORCA Flash4.0 CMOS camera (Hamamatsu Photonics, Japan). 45,000-50,000 frames were collected at 100 fps. Ca_V_1.2 cluster areas and events per pixel were analyzed from 10 nm localization maps using Fiji:ImageJ (NIH).

### Immunostaining and Airyscan microscopy

Isolated myocytes were plated on poly-L-lysine/laminin-coated coverslips (37 °C, 45 min), then fixed in 4 % paraformaldehyde in PEM buffer (80 mM <u>P</u>IPES, 5 mM <u>E</u>GTA, 2 mM <u>M</u>gCl_2_, 37 °C, 10 min). After permeabilization (0.5% Triton-X 100, 10 min) and blocking (50% SEA Block, 0.5% Triton-X100 in PBS, 1 hr), samples were immunolabeled overnight at 4 °C with rabbit polyclonal IgG anti-Ca_V_1.2 (1:300; Alomone labs ACC-003) and one of the following: mouse monoclonal anti-EEA1 IgG1 (1:250; BD Bioscience 610456), mouse monoclonal IgG1 anti-Rab11a (sc-166523, 1:250; Santa Cruz Biotechnology Inc., Dallas, TX, USA), or mouse monoclonal IgG1 anti-Rab7 (1:250; Santa Cruz sc-376362) antibodies. Following PBS washes, samples were labeled with Alexa Fluor 488 anti-rabbit and Alexa Fluor 647 anti-mouse secondary antibodies (1:1000, 1 hr). Images were acquired using a Zeiss LSM 880 Airyscan microscope with 63×/1.40 oil objective and Zen Software (Carl Zeiss Microscopy, LLC., White Plains, NY, USA).

### Whole-cell patch clamp electrophysiology

Ca_V_1.2 currents (*I*_Ca_) were recorded using borosilicate pipettes (1-3 MΩ) filled with internal solution (87 mM Cs-aspartate, 20 mM CsCl, 1 mM MgCl_2_, 10 mM HEPES, 10 mM EGTA, 5 mM MgATP; pH 7.2). Myocytes were initially perfused with Tyrode’s solution (140 mM NaCl, 5 mM KCl, 10 mM HEPES, 10 mM glucose, 1 mM MgCl_2_, 2 mM CaCl_2_; pH 7.4), then switched to recording solution (5 mM CsCl, 10 mM HEPES, 10 mM glucose, 140 mM NMDG, 1 mM MgCl_2_, 2 mM CaCl_2_; pH 7.3) after whole-cell configuration was achieved. To minimize rundown, cells were held at −80 mV for 5 min before recording. The voltage protocol stepped to −40 mV (100 ms) to inactivate Na^+^ channels, followed by 300 ms steps from −60 to +90 mV. Currents were sampled at 10 kHz, filtered at 2 kHz (Axopatch 200B), and digitized (Digidata 1550B; Molecular Devices, Sunnyvale, CA, USA). Analysis used Clampfit software with liquid junction potential correction (−10 mV). Current-voltage relationships were fit with Boltzmann equations (GraphPad Software Inc., La Jolla, CA, USA).

### Calcium transient recordings

Myocytes were loaded with Fluo-4 AM (Thermo Fisher Scientific, 20 min in the dark per manufacturers protocol), centrifuged (300 rpm, 2 min) to allow aspiration of the excess dye, then resuspended in Tyrode’s solution for de-esterification (20 min). Line-scan confocal microscopy (Zeiss LSM 880, 63×/1.40 objective, 488-nm excitation) was used to record Ca^2+^ transients from cells paced to steady state (Myopacer, IonOptix, LLC., Westwood, MA; 12-15V, 1 Hz) during continuous perfusion with Tyrode’s. Fluorescent signals were converted to [Ca^2+^]_i_ using the pseudo-ratiometric approach^33^ and the equation: [Ca^2+^]_i_= K_d_(F/F_0_)/(K_d_/[Ca^2+^]_i-rest_+1−F/F_0_) as previously described^34, 35^.

### Preparation of whole-heart lysates

Hearts from infused mice were snap-frozen in liquid nitrogen and homogenized in RIPA buffer (R26200, Research Products International, Mount Prospect, IL USA) containing protease inhibitor, microcystin (Insolution microcystin-LR, Thermo Fisher Scientific, 4 μM), and sodium fluoride (1 mM) using a Bullet Blender Storm 24 centrifuge (Next Advance; Troy, NY, USA, 4 °C, 20 min). Protein concentration in the extracted supernatant was determined by BCA assay (Prometheus Biosciences, Rahway, New Jersey, USA).

### Western Blot

Lysate samples (20 μg) were denatured in SDS buffer (Invitrogen, Waltham, MA, USA; 70 °C for at least 10 min), separated by SDS-PAGE on 4-12 % acrylamide gradient gels, and transferred to PVDF membranes (Bio-Rad Laboratories, Hercules, CA, USA) in transfer buffer (95.9 mM glycine, 12.5 mM Tris, 150 ml MeOH, up to 1 L H_2_0; 50 V, 10 hrs, 4 °C). After Ponceau staining (Research Products International, Mount Prospect, IL, USA), membranes were blocked for 1 hr in 5 % milk in a TBST solution (150 mM NaCl, 10 mM Tris-HCl, pH 7.4 with 0.2 % Tween) and incubated with primary antibodies (4 °C, overnight): Ca_V_1.2 (NeuroMab anti-Ca_V_1.2 II-III, 1:200 or Alomone Labs, Jerusalem, Israel, ACC-003, 1:300), pSer1928 (NeuroMab L132/68.1, 1:10), PKA catalytic-α (Santa Cruz sc28315, 1:500 in 2 % BSA), RyR2-pSer2814 (Badrilla, Leeds, UK, A010-31, 1:500), RyR2-pSer2808 (Badrilla, A010-30, 1:500), RyR2 (Alomone Labs, ARR-002, 1:200), GAPDH (Proteintech, Rosemont, IL, 1:1000), and α-tubulin (Invitrogen, MA1-80017, 1:1000). After TBST washes, membranes were incubated for 1 hr with HRP-conjugated secondary antibodies (Bio-Rad/Jackson ImmunoResearch 211-032-171; Biorad 170516; or Biorad 5204-2504; all diluted 1:10,000), washed, and developed using chemiluminescence reagents (Immobilon Classico, Sigma-Aldrich; Prometheus ProSignal Femto, Genesee Scientific, El Cajon, CA, USA) on autoradiography film (Amersham Hyperfilm ECL, Cytivia, Global Life Sciences Solutions USA, LLC., Marlborough, MA, USA). Multiple exposures ensured linear-range signals. Band intensities were quantified by densitometry in Adobe Photoshop, normalizing phospho-proteins to total protein and the average of protein loading using Ponceau/loading controls.

### Lipid Mass Spectrometry

Phosphoinositide quantification from whole heart samples was performed using ultra-high pressure liquid chromatography coupled to tandem mass spectrometry (UPLC-MS/MS) as previously described^36, 37^. Endogenous lipids and internal standards were extracted using N-butanol/chloroform, methylated, then separated on a C4 column using an acetonitrile/formic acid gradient. The column eluate was infused with sodium formate and analyzed using a Waters XEVO TQ-S mass spectrometer in multiple reaction monitoring mode (MRM) with electrospray ionization and positive ion mode. Peak areas were quantified using MassLynx software, normalized to synthetic standards, and corrected for total protein content.

### T-tubule organization analysis

Freshly isolated myocytes were stained with the membrane dye di-8-ANEPPS (Thermo Fisher) for 30 minutes in the dark. After centrifugation (300 rpm, 2 min) and resuspension in Tyrode’s solution, cells were imaged using a Zeiss LSM 880 Airyscan super-resolution microscope with a 63×/1.40 oil objective. T-tubule organization was quantified from the acquired images using ImageJ/Fiji with the TTorg plugin^38^.

### Echocardiography and Doppler Imaging

Cardiac structure and function were evaluated in the same animals before and after one week of infusion using a Vevo 2100 ultrasound system equipped with a MS 550D probe (22-55 MHz, VisualSonics, Fujifilm, Toronto, ON, Canada). Two-dimensional M-mode echocardiography and pulsed-wave Doppler measurements were performed on isoflurane-anesthetized (2%) mice to assess systolic and diastolic function. Measured parameters included: left ventricular anterior and posterior wall dimensions (LVAW, LVPW) and inner diameter (LVID) in both diastole and systole, fractional shortening, LV mass, heart rate, mitral valve E/A ratio, and isovolumetric relaxation time (IVRT).

### Statistical Analysis

Data are presented as mean ± SEM, with *N* indicating number of animals and *n* indicating number of cells. Statistical comparisons were performed using GraphPad Prism software with paired or unpaired Student’s t-tests for two-group comparisons, and two-way ANOVA with appropriate post-hoc tests for multiple group comparisons. Data that were not normally distributed were analyzed with non-parametric tests. Differences were considered significant at *P* < 0.05.

## Results

### Whole-heart phospholipid species’ levels are altered by chronic AngII treatment

Given our prior finding that acute treatment of mouse hearts with AngII (100 nM for 5 mins via Langendorff perfusion) significantly decreased PIP_2_ (∼44 % reduction) and PIP (∼63 % reduction) levels in Langendorff-perfused hearts, we began this investigation by asking if a similar perturbation of phosphoinositide levels occurs upon chronic *in vivo* infusion of AngII. To do this, we performed an exquisitely sensitive interrogation of the cardiac phospholipid landscape by performing phospholipid mass spectrometry on whole hearts from mice subjected to a week-long chronic infusion of AngII (1 mg/kg/day) or saline (same volume vehicle control) via subcutaneous osmotic mini-pump. Figure 1A outlines a simplified view of phosphoinositide metabolism, showing the lipid kinases and phosphatases that catalyze the inter-conversion of the various phosphoinositide species. Our results indicate an extensive alteration of all quantified phosphoinositide species in the AngII-infused group, indicating pervasive phospholipid imbalance. Specifically, we measured a 37.30 ± 12.15 % decrease in PI, a 38.91 ± 13.80 % increase in PIP, a 43.71 ± 14.56 % decrease in PIP_2_, and a 35.74 ± 14.32 % decrease in PIP_3_ in AngII-infused hearts compared to saline-infused controls (Figure 1B-E). We predict that such a profound imbalance of these phosphoinositide species will have repercussions for the activity of the myriad of cardiac ion channels, exchangers, organelles, and cellular processes that are regulated by these lipids.

**Figure 1.**
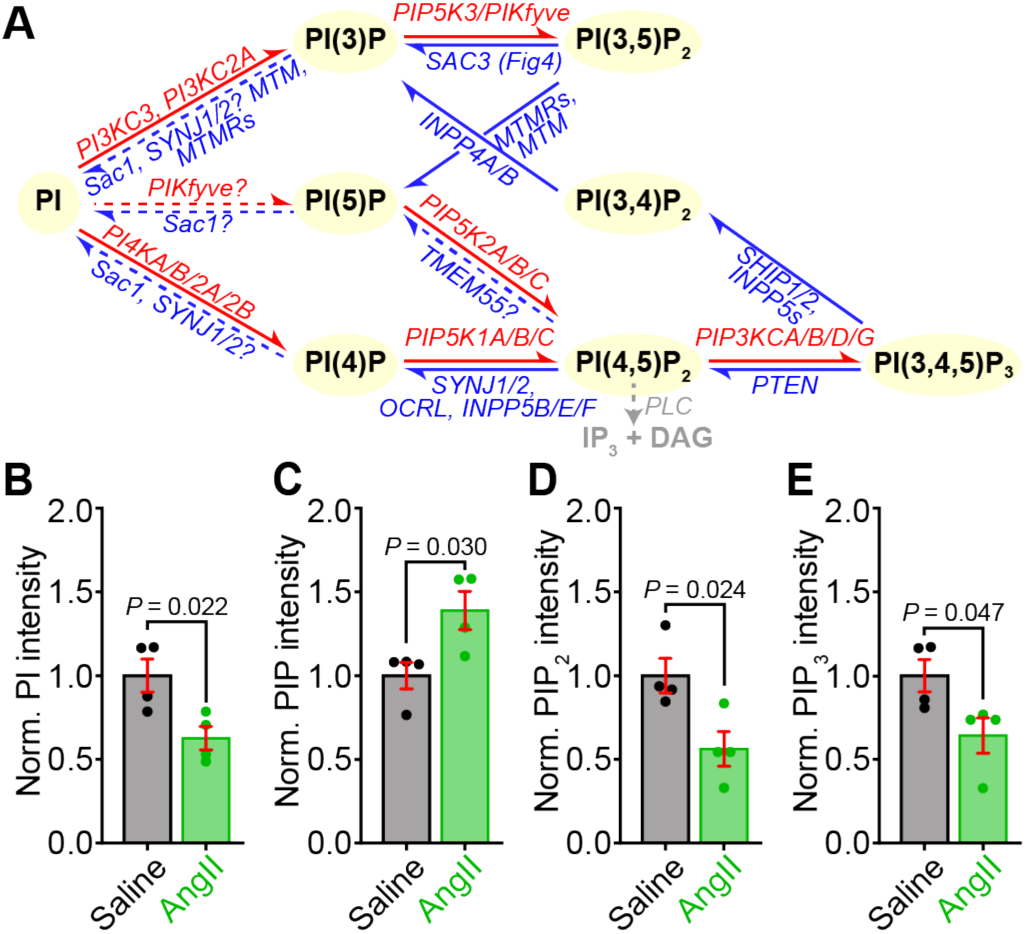
Phospholipid mass spectrometry reveals whole-heart phospholipid imbalances in chronically AngII-stimulated mice. **A,** The homeostatic phospholipid metabolism pathway. **B-E**, Bar plots summarizing UPLC-MS/MS measurements of PI (**B**), PIP (**C**), PIP_2_ (**D**), and PIP_3_ (**E**) in whole heart lysates from saline or Ang II-infused mice, (*N* = 4 for each). Data were analyzed using unpaired, two-tailed Student’s *t*-tests. Error bars indicate SEM.

### Chronic AngII infusion reduces Ca_V_1.2 cluster and expression

Having established that chronic AngII-infusion *in vivo* results in a significant decrease in PIP_2_, we proceeded with an examination of Ca_V_1.2 channel localization and clustering in ventricular myocytes isolated from the hearts of mice subjected to chronic infusion of AngII or saline. This was a logical starting point for the current investigation given our previous finding that acute AngII exposure (100 nM, 5 min) selectively reduced t-tubular Ca_V_1.2 clusters by ∼30% compared to controls while sparing the crest population^29^. Accordingly, cells were fixed, immunostained against the α_1C_-subunit of Ca_V_1.2, and imaged using super-resolution single-molecule localization microscopy (SMLM). Analysis revealed t-tubule localized Ca_V_1.2 channel cluster areas from AngII-infused mice were 23.5 ± 8.2 % smaller on average than those in saline-infused controls (Figure 2A-B). Chronic AngII was also associated with a 54.3 ± 20.5 % decrease in the number of detectable events per 10 nm pixel in single-molecule localization maps (Figure 2C). This measure provides an indication of the total channel expression level in the imaged area of the cell. We previously reported a similar (47.9 %) decrease in this metric in myocytes subjected to acute AngII treatment^29^. Examination of sarcolemmal crest localized Ca_V_1.2 channels revealed no significant impact of chronic AngII-infusion on Ca_V_1.2 channel expression or clustering (Figure 2D-F). Taken together, these SMLM data suggest that chronic AngII-infusion reduced the t-tubule sarcolemma population of Ca_V_1.2 channels in ventricular myocytes.

**Figure 2.**
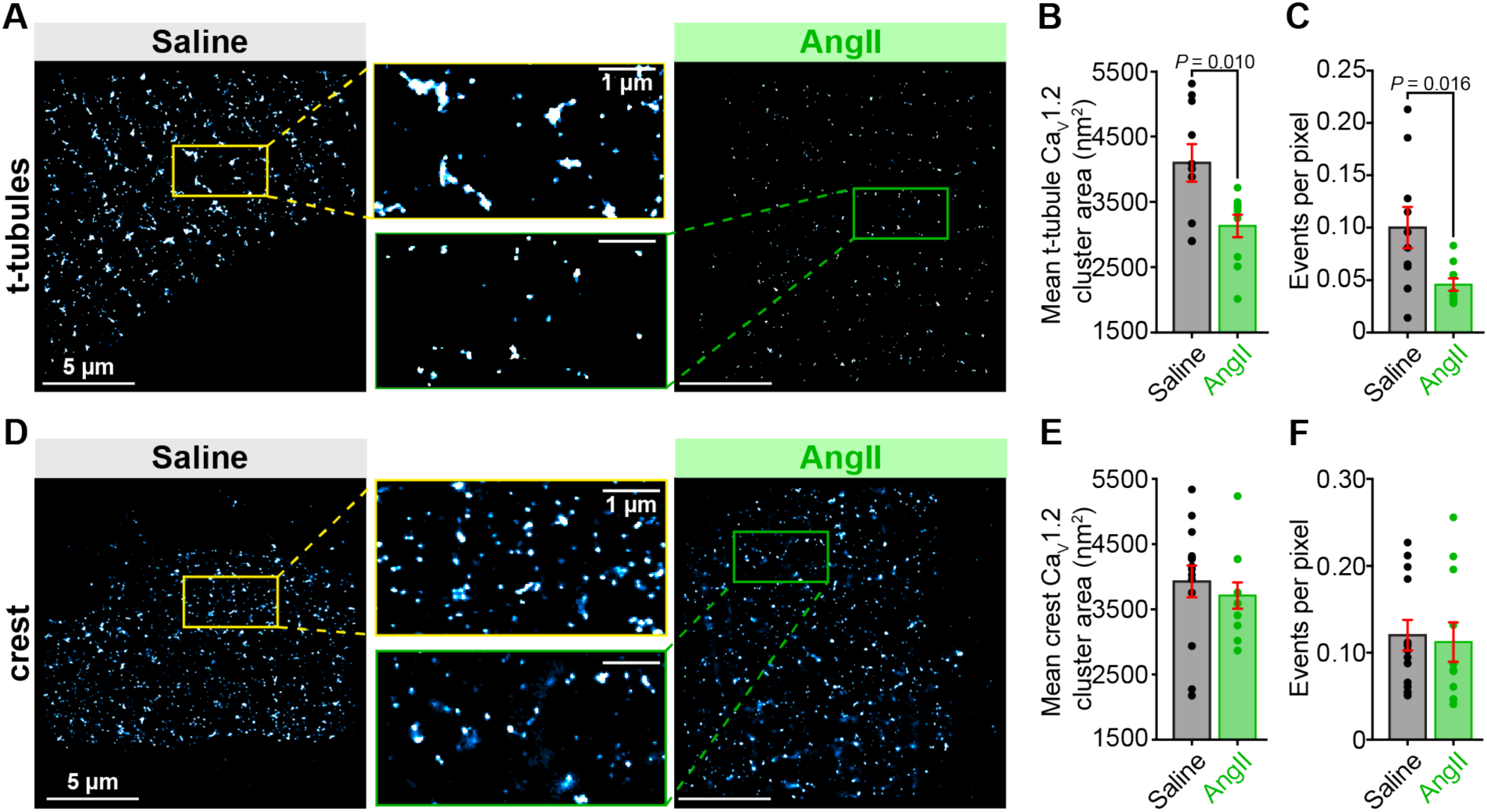
Chronic AngII-infusion reduces t-tubular, but not crest-localized Ca_V_1.2 cluster size and expression. SMLM localization maps of fixed ventricular myocytes immunostained to examine Ca_V_1.2 channel distribution on the t-tubules (**A**) (saline *N* = 3, *n* = 10, AngII *N* = 3, *n* = 10) and sarcolemmal crest (**D**) (saline *N* = 3, *n* = 13, AngII *N* = 3, *n* = 11) under saline control (*left*) or chronically AngII-stimulated (*right*) conditions. Aligned dot plots showing mean Ca_V_1.2 channel cluster area (**B, E**), and the events per pixel (**C, F**) in each location and condition. Unpaired Student’s t-test, \**P* < 0.05. Error bars indicate SEM.

### Chronic AngII increases Ca_V_1.2 channel localization to endosomes

We next investigated the fate of the Ca_V_1.2 channels that were absent from the t-tubule sarcolemma. If the channels were endocytosed into the endosomal recycling system, then a testable prediction is that endosomal reservoirs of Ca_V_1.2 channels should be enriched within cardiomyocytes isolated from chronic AngII-infused mice compared to those from saline-infused mice. To test this hypothesis, we performed Airyscan super-resolution imaging on myocytes isolated from chronically saline-infused control and AngII-infused mice. Cells were fixed and immuno-stained against Ca_V_1.2 and various endosome populations including EEA1-positive early endosomes, Rab11a-positive recycling endosomes, and Rab7-positive late endosomes. Chronic AngII treatment resulted in an 18.76 ± 6.23 % increase in EEA1/Ca_V_1.2 colocalization (Figure 3A-B), a 38.97 ± 11.26 % increase in Rab11a/Ca_V_1.2 colocalization (Figure 3C-D), and a 19.53 ± 5.04 % increase in Rab7/Ca_V_1.2 colocalization (Figure 3E-F), compared to saline-infused controls. These results suggest that one week of chronic AngII-infusion triggers endocytosis of Ca_V_1.2 channels from the t-tubule sarcolemma and subsequent collection on endosomal reservoirs within early, recycling, and late endosomes.

**Figure 3.**
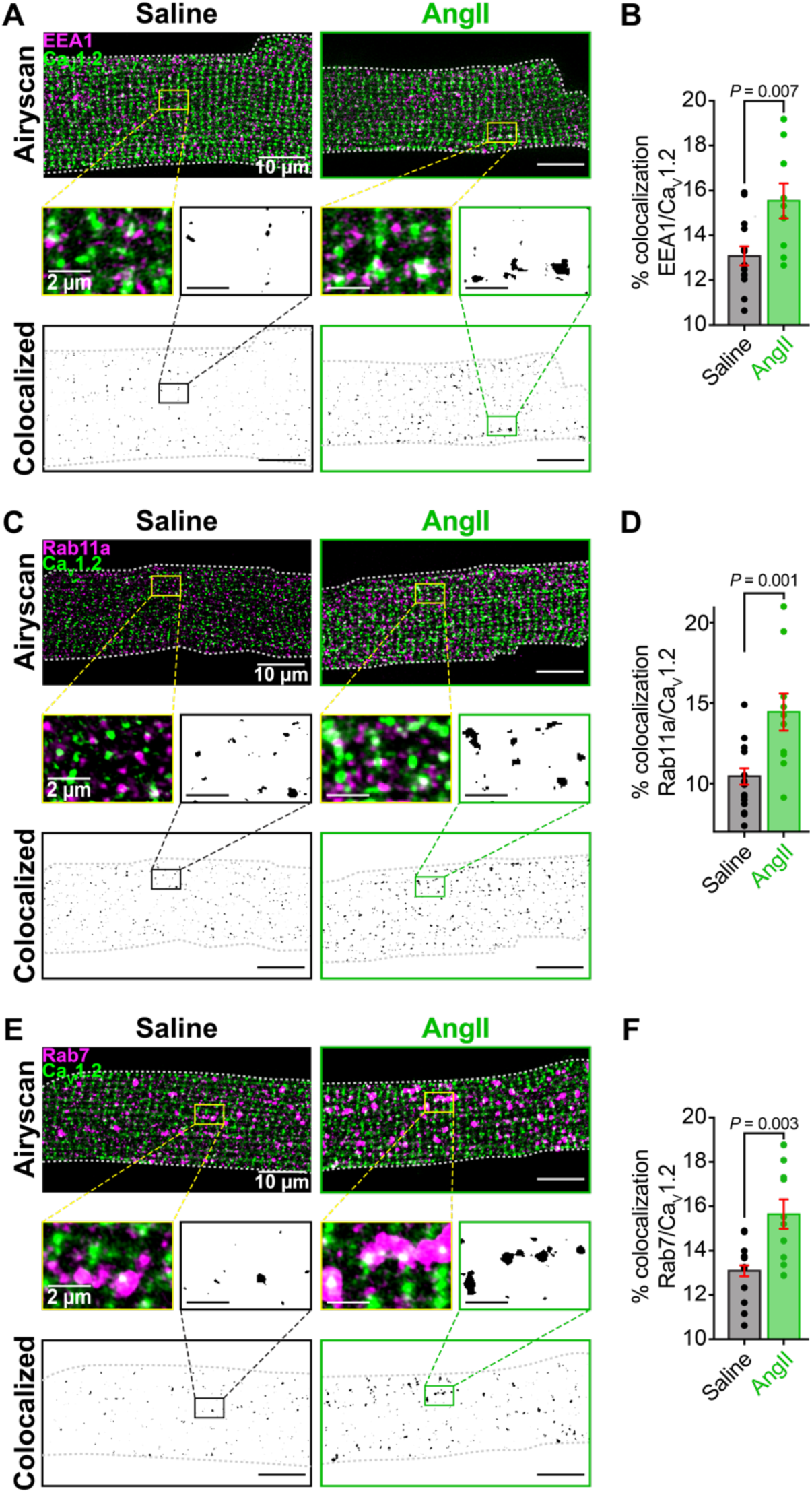
Chronic AngII stimulation induces endocytosis of Ca_V_1.2 channels into early, recycling, and late endosomes. **A**, Two-color Airyscan super-resolution images showing distributions of Ca_V_1.2 and EEA1^+^ early endosomes in representative Saline (*left*) and AngII-stimulated (*right*) myocytes. Binary colocalization maps (*bottom*) display pixels in which Ca_V_1.2 and endosomal expression fully overlap. **B**, Bar plot summarizing percent colocalization between EEA1 and Ca_V_1.2 in control (*N* = 4, *n* = 14) and AngII conditions (*N* = 3, *n* = 9). **C**, Immunostaining of Ca_V_1.2 and Rab11a recycling endosomes. **D**, Bar plot summarizing results from Saline (*N* = 4, *n* = 16) and AngII-stimulated (*N* = 4, *n* = 10) myocytes. **E,** Immunostaining of Ca_V_1.2 and Rab7^+^ late endosomes. **F,** Bar plot summarizing results from Saline (*N* = 4, *n* = 12) and AngII-stimulated (*N* = 3, *n* = 10) myocytes. Unpaired Student’s *t*-test, \**P* < 0.05. Error bars indicate SEM.

### I_Ca_ is preserved with chronic AngII-infusion

A decrease in Ca_V_1.2 channel clustering and expression at the t-tubules would be predicted to reduce whole-cell Ca^2+^-currents (*I*_Ca_) in myocytes from the chronic AngII-infused mice. We tested this using whole-cell patch clamp electrophysiology. As anticipated, chronic AngII-infusion was accompanied by an increase in heart weight/body weight (HW/BW) ratio, implying cardiac hypertrophy, and enhanced ventricular myocyte capacitance, indicating cellular hypertrophy (Figure 4A-B). However, contrary to our prediction, and despite the apparent reduction in Ca_V_1.2 channel expression, *I*_Ca_ density was not significantly altered by chronic AngII-infusion compared to saline-infused controls (Figure 4C-E). With fewer channels in the sarcolemma, as indicated by reduced events/pixel in Figure 2C, one hypothesis to explain the preservation of *I*_Ca_ density is a functional upregulation of the remaining channels. A well-studied mechanism to increase Ca_V_1.2 channel activity in the heart occurs via *β*-adrenergic receptor stimulation and downstream signaling through adenylyl cyclase, leading to cAMP production and increased PKA-mediated phosphorylation of Ca_V_1.2 channel complexes. PKA-phosphorylated Ca_V_1.2 channels exhibit two distinctive characteristics: First, they show a left-ward shift in their voltage dependence of conductance (G/G_max_), enabling channel opening at more negative membrane potentials and thus requiring weaker depolarization for activation. Consistent with this, our analysis of Ca_V_1.2 biophysical characteristics in myocytes from AngII-infused mice revealed a significant left-ward shift in the G/G_max_ and a more hyperpolarized voltage of half-maximal activation (V_1/2_) compared to saline controls (Figure 4F-G). Second, *β*-adrenergic regulation modifies inactivation kinetics, causing slower voltage-dependent inactivation (VDI) while accelerating calcium-dependent inactivation (CDI)^39–43^. When comparing AngII-infused cells to saline-infused controls generating similar magnitude *I*_Ca_ (and thus presumably similar CDI), we observed significantly slower overall channel inactivation in the AngII-infused cells (Figure 4C and H). Together, these data indicate preservation of ventricular myocyte *I*_Ca_ in AngII-infused mice may occur due to a compensatory sympathetic response.

**Figure 4.**
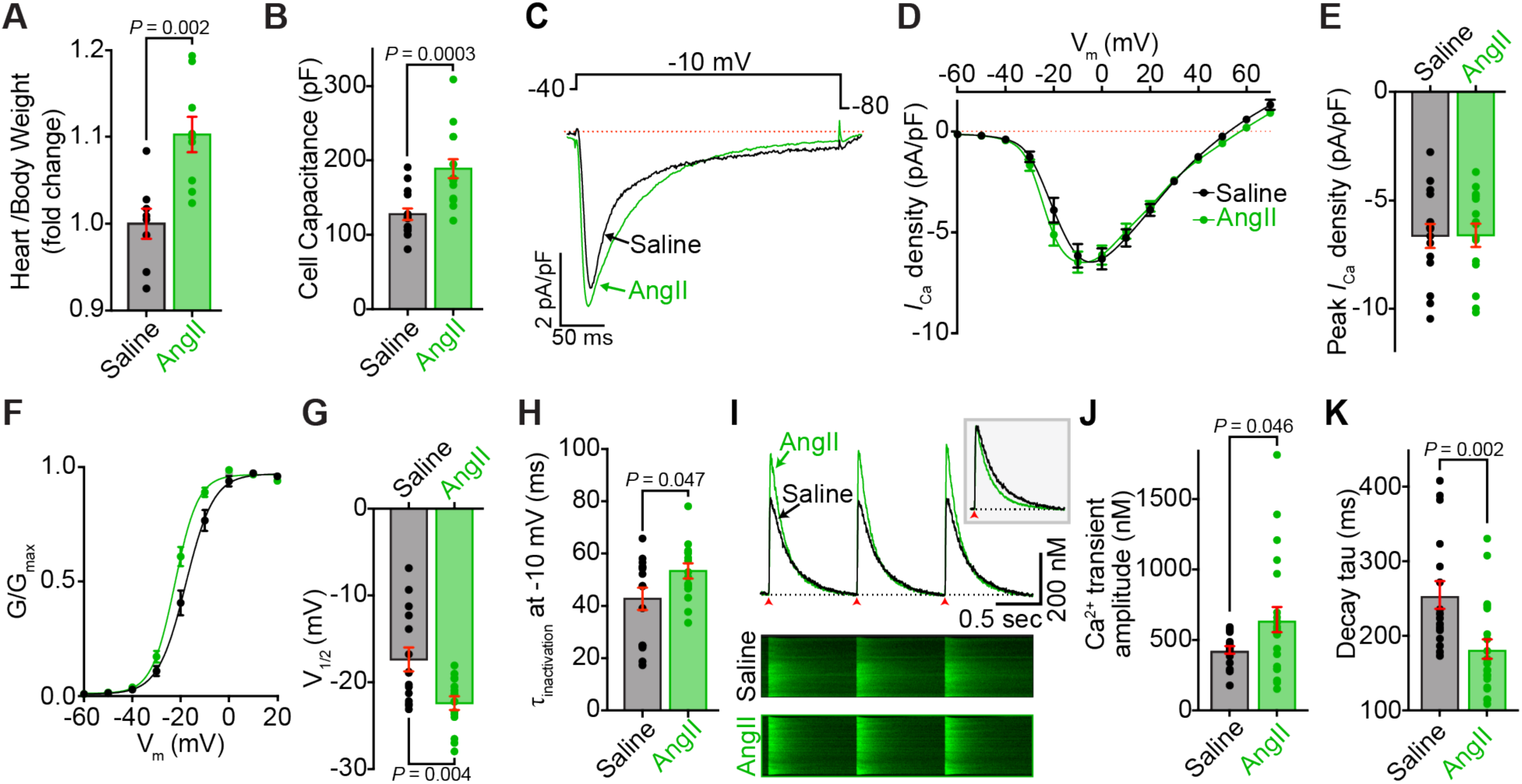
Functional analysis reveals conserved *I_Ca_* and augmented Ca^2+^ transients in cardiomyocytes, and hypertrophy in chronic AngII-infused hearts. **A,** Heart weight to body weight ratio of saline-infused (*N* = 8) and AngII-infused (*N* = 9) mice. **B,** Cell capacitance of ventricular myocytes isolated from saline and AngII-infused hearts. **C,** Representative whole-cell *I*_Ca_ elicited from isolated ventricular myocytes from saline-infused (black), and AngII-infused (green) mouse hearts, during a 300 ms depolarization step to −10 mV **D**, Current-voltage relationships of *I*_Ca_ density in myocytes subjected to test potentials from −60 mV to +70 mV. **E,** Peak *I*_Ca_ density of myocytes from saline (*N* = 6, *n* = 15), and AngII (*N* = 5, *n* = 15) groups. **F**, Voltage-dependence of normalized conductance (G/G_max_) with Boltzmann fits, and **G**, the associated half-maximal activation voltages (V_1/2_). **H**, Time constant of *I*_Ca_ inactivation at −10 mV from saline and AngII groups. **I,** Representative EFS-evoked Ca^2+^ transients (*top*) and corresponding line-scan images (*bottom*) from fluo4-AM loaded myocytes from saline (black) and AngII-infused (green) hearts. Arrows indicate 1 Hz stimulation. **J**, Fold-change in Ca^2+^ transient amplitude, and **K,** decay tau of Ca^2+^ transients (saline: *N* = 3, *n* = 17, AngII: *N* = 3, *n* = 22). Data were analyzed using unpaired Student’s t-tests. Error bars indicate SEM.

### Ca^2+^ transient amplitude and rate of decay is enhanced with chronic AngII-infusion

To investigate the functional impact of chronic AngII infusion on excitation-contraction (EC) coupling, we analyzed intracellular Ca^2+^ transients. We previously reported that acute treatment of isolated myocytes with AngII caused a significant (∼16 %) reduction in Ca^2+^ transient amplitude in isolated ventricular myocytes that occurred due to reduced trigger Ca^2+^ influx downstream of AT_1_R/G_q_/PLC catalyzed PIP_2_-hydrolysis and consequent endocytosis of a proportion of the sarcolemmal Ca_V_1.2 channel population. As discussed above, in myocytes isolated from chronically AngII-perfused hearts *I*_Ca_ amplitude was preserved despite reduced clustering and sarcolemmal localization of Ca_V_1.2. In line with that, we did not observe a reduction in Ca^2+^transient amplitudes in myocytes from chronically AngII-infused mice. On the contrary, we observed an *increase* in Ca^2+^ transient amplitude (Figure 4I-J) compared to saline-infused controls. A significant acceleration in Ca^2+^ transient decay kinetics was also observed in chronic AngII cells versus saline controls (Figure 4I; *inset*, and K), indicating an enhanced rate of store-refilling. Increased Ca^2+^ transient amplitude and accelerated rates of store refilling are both features of *β*-adrenergic regulation of cardiac function. Together with the patch clamp results, these data suggest that chronic AngII-infusion triggers enhanced sympathetic outflow and *β*-adrenergic signaling.

### Ca_V_1.2 and RyR2 phosphorylation is enhanced after chronic AngII

With several data sets pointing toward *β*-adrenergic stimulation in chronic AngII-infused mice, we next examined the expression and phosphorylation state of Ca_V_1.2 and RyR2 in a series of western blots performed on whole heart lysates obtained from saline-infused and chronically AngII-infused mice. No appreciable changes in total Ca_V_1.2α_1c_ expression were observed between the two groups (Figure 5A-B). However, phosphorylated Ser1928-Ca_V_1.2 was significantly increased in heart lysates from chronic AngII-infused mice compared to saline controls (Figure 5C). Stimulation of *β*-adrenergic receptors in cardiomyocytes produces substantial PKA-mediated phosphorylation of the S1928 residue on Ca_V_1.2^44^. Although S1928 phosphorylation is not essential for *β*-adrenergic-mediated potentiation of *I*_Ca_ ^45, 46^, detection of enhanced phospho-S1928 levels is still an accurate indicator of enhanced PKA activity. Examination of the expression of the PKA catalytic subunit revealed an increase in its abundance in AngII-infused mouse hearts compared to saline controls, which may also contribute to the enhanced PKA-mediated phosphorylation of Ca_V_1.2 channel complexes (Figure 5D-E). It is well known that cAMP/PKA signaling desensitizes and gives way to CaMKII signaling during chronic *β*-adrenergic signaling^47^; if AngII is triggering chronic *β*-adrenergic signaling, then it is possible that CaMKII activity is also enhanced. To our knowledge, at time of writing, there are no validated commercial antibodies that can reliably detect CaMKII-mediated phosphorylation of Ca_V_1.2; however, there are well-validated antibodies for the CaMKII (S2814) and PKA (S2808) phospho-sites on RyR2. We thus examined the extent of PKA and CaMKII-mediated phosphorylation of RyR2. Total RyR2 abundance was not altered (Figure 5F-G and I); however, both CaMKII-mediated phosphorylation of S2814 (Figure 5F and H) and PKA-mediated phosphorylation of S2808 (Figure 5I-J) was significantly increased in lysates from chronic AngII-infused mice compared to saline-infused controls. These results imply that chronic AngII-infusion leads to an increase in CaMKII and PKA activity in the heart, fueling enhanced phospho-regulation of their protein targets, including Ca_V_1.2, RyR2, and phospholamban.

**Figure 5.**
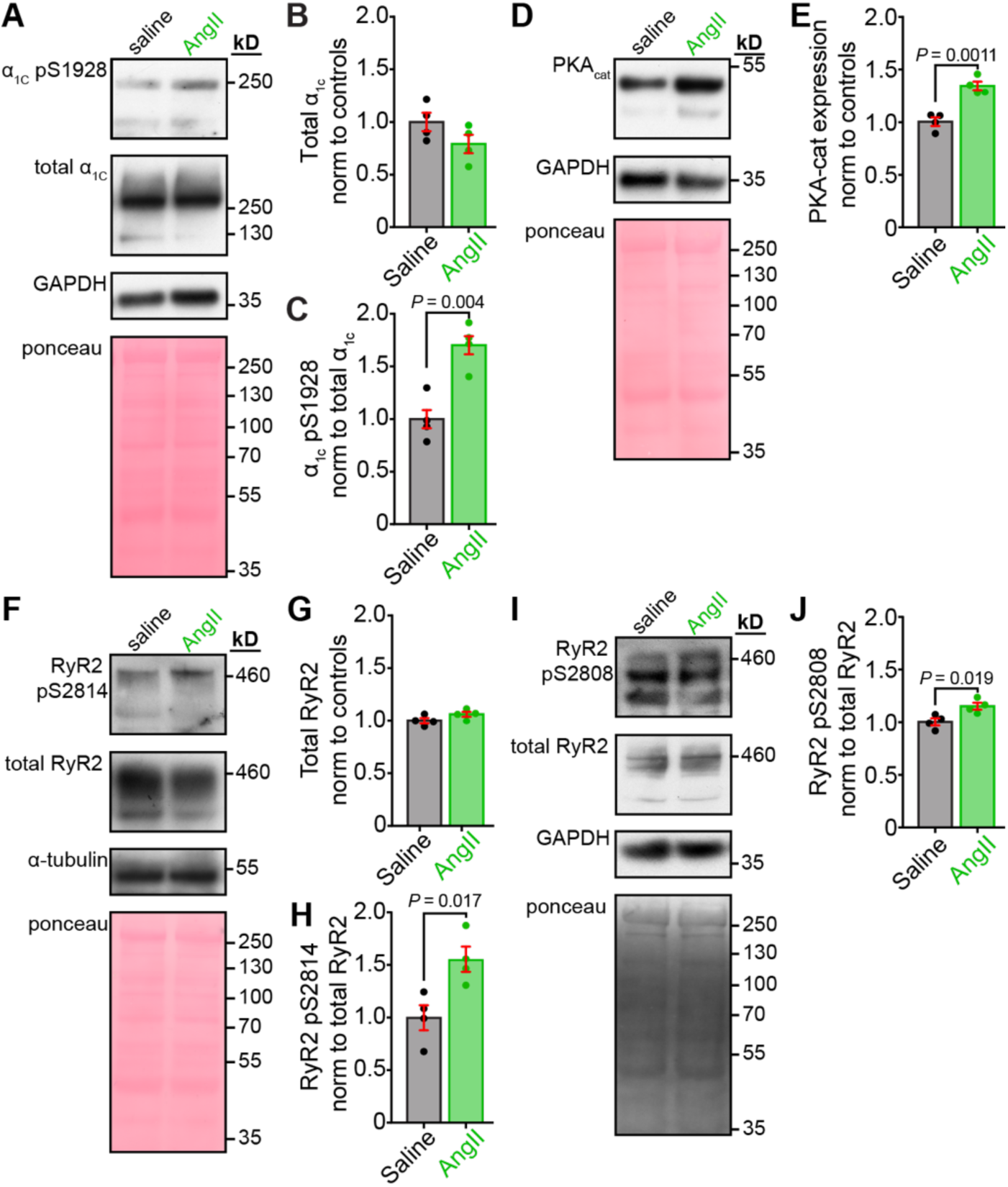
Chronic AngII infusion enhances PKA- and CaMKII-mediated phosphorylation of Ca_V_1.2 and RyR2. Representative western blots and quantification from saline and AngII-infused heart lysates (*N* = 4/group). **A**, Immunoblots showing Ca_V_1.2 pSer1928, total Ca_V_1.2α_1C_, with GAPDH and Ponceau loading controls. Quantification of **B**, Ca_V_1.2 pSer1928 and **C**, total Ca_V_1.2α_1C_ expression. **D**, Immunoblots of PKA catalytic α with loading controls and **E**, corresponding quantification. **F**, Immunoblots showing RyR2 pS2814, total RyR2, with α-tubulin and Ponceau controls. Quantification of **G**, RyR2 pS2814 and **H**, total RyR2. **I**, Immunoblots of RyR2 pS2808 with corresponding total RyR2, GAPDH, and Ponceau controls, and **J**, quantification of RyR2 pS2808. Data were analyzed using unpaired Student’s t-tests. Error bars indicate SEM.

### Chronic AngII induces hypertrophic remodeling that impairs cardiac function

Chronic infusion of AngII leads to hypertension due to its potent vasopressor effect^48^. Cardiac hypertrophy can develop as a compensatory mechanism to maintain cardiac output in response to the chronic increase in afterload caused by hypertension. While this adaptive response is initially beneficial, the growth of the ventricular wall can become pathogenic, leading to impaired relaxation and diastolic dysfunction, which can progress to HF. Accordingly, we performed echocardiography and doppler imaging to assess systolic and diastolic function in paired experiments where cardiac function was measured prior to implantation of the mini-osmotic pump and again after the one-week infusion with either AngII or saline. Consistent with the capacitance and gross HW/BW results in Figure 4A and B, left ventricular wall thickness and overall left ventricular (LV) mass were increased by the AngII infusion compared to saline controls (Figure 6A-E), confirming cardiac hypertrophy. We also observed that the inner diameter of the LV chamber (LVID) was significantly reduced (Figure 6F), suggesting that the hypertrophic growth of the wall had encroached on the LV cavity. This chamber constriction in diastole would be predicted to limit end-diastolic volume (EDV), and indeed, that is what we observed (Figure 6G), along with a decreased end-systolic volume (which did not reach significance), stroke volume, and cardiac output (Figure 6H, I, J). One may have predicted an increase in heart rate due to the sympatho-excitatory effects of AngII, but this was not evident in our recordings (Figure 6K). Fractional shortening was also significantly reduced in the AngII-infused mice (Figure 6L) despite the maintained *I*_Ca_ and larger Ca^2+^ transients in individual ventricular myocytes (see Figure 4). This discrepancy suggests the presence of pathological remodeling/fibrosis, which likely restricts contractility and limits cardiac output.

**Figure 6.**
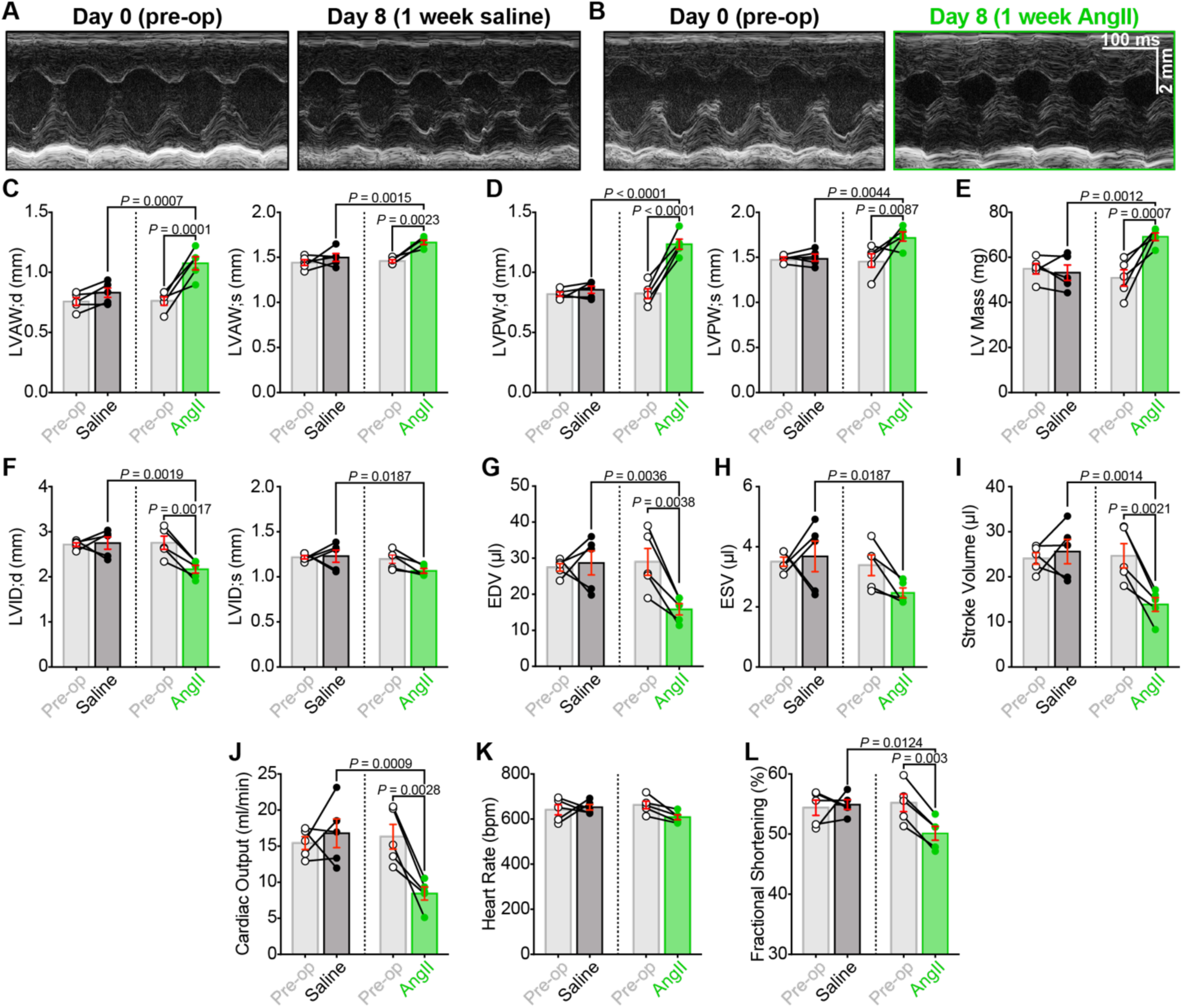
Chronic AngII-infusion alters cardiac structure and impairs systolic function. Representative short-axis M-mode echocardiograms of hearts from **A,** saline and **B,** AngII-infused mice before (*left*) and after 1 week of infusion (*right*), recorded from the same mice before (pre-op, *left*) and after 1 week of infusion (post-infusion, *right*). **C-L,** Bar plots comparing paired echocardiographic parameters between pre-op and post-infusion timepoints in saline (*N* = 5) and AngII-infused (*N* = 5) mice, including: **C**, left ventricular anterior wall thickness and **D,** posterior wall thickness measured in diastole (*left*) and systole (*right*); **E,** left ventricular mass (LV mass); **F,** left ventricular inner diameter (LVID); **G,** end diastolic volume (EDV); **H,** end systolic volume (ESV); **I,** stroke volume; **J,** cardiac output; **K,** heart rate, and **L,** fractional shortening. Data analyzed using two-way ANOVA with Tukey’s post-hoc tests; error bars indicate SEM.

### Chronic AngII infusion results in diastolic dysfunction

The changes in EDV and LV chamber diameter hint at impaired relaxation and possible diastolic dysfunction; thus, we performed Doppler imaging to examine blood flow through the mitral valve. At the conclusion of systole, as diastole begins, blood flow from the left atrium (LA) to the left ventricle is initially a passive process as the pressure in the relaxing and relatively empty LV drops below the pressure in the LA. The pressure differential allows the mitral valve to open and passive blood flow down the pressure gradient. This passive blood flow can be observed using pulsed-wave Doppler imaging and appears as the so-called E-wave. The A wave reflects active blood flow during the subsequent contraction of the LA, which provides the “atrial kick.” Both E and A waves are visible and labeled on Figure 7 A-B. E wave amplitude was significantly reduced after AngII-infusion compared to the same mice pre-op (before the osmotic minipump was implanted) or saline-infused controls (Figure 7C). This indicates less passive filling of the LV and suggests an altered pressure gradient between the LA and the LV, which may occur due to decreased LV compliance and impaired diastolic relaxation. If the LV relaxes more slowly or incompletely, LV pressure will be higher than normal, thus reducing the LA-LV pressure differential during the passive filling phase. AngII-infusion did not significantly alter A wave amplitude; thus, a significantly reduced E/A ratio of the transmitral flow was observed (Figure 7D). We also observed a prolongation of the isovolumetric relaxation time (IVRT, Figure 7E). This reflects the time between aortic valve closure and mitral valve opening during isovolumic relaxation of the LV. The mitral valve only opens when LA pressure exceeds that of the LV. Thus, the IVRT prolongation suggests a less steep drop in LV pressure during relaxation, i.e., impaired diastolic relaxation. Together, these data indicate that chronic AngII-infusion produces diastolic dysfunction.

**Figure 7.**
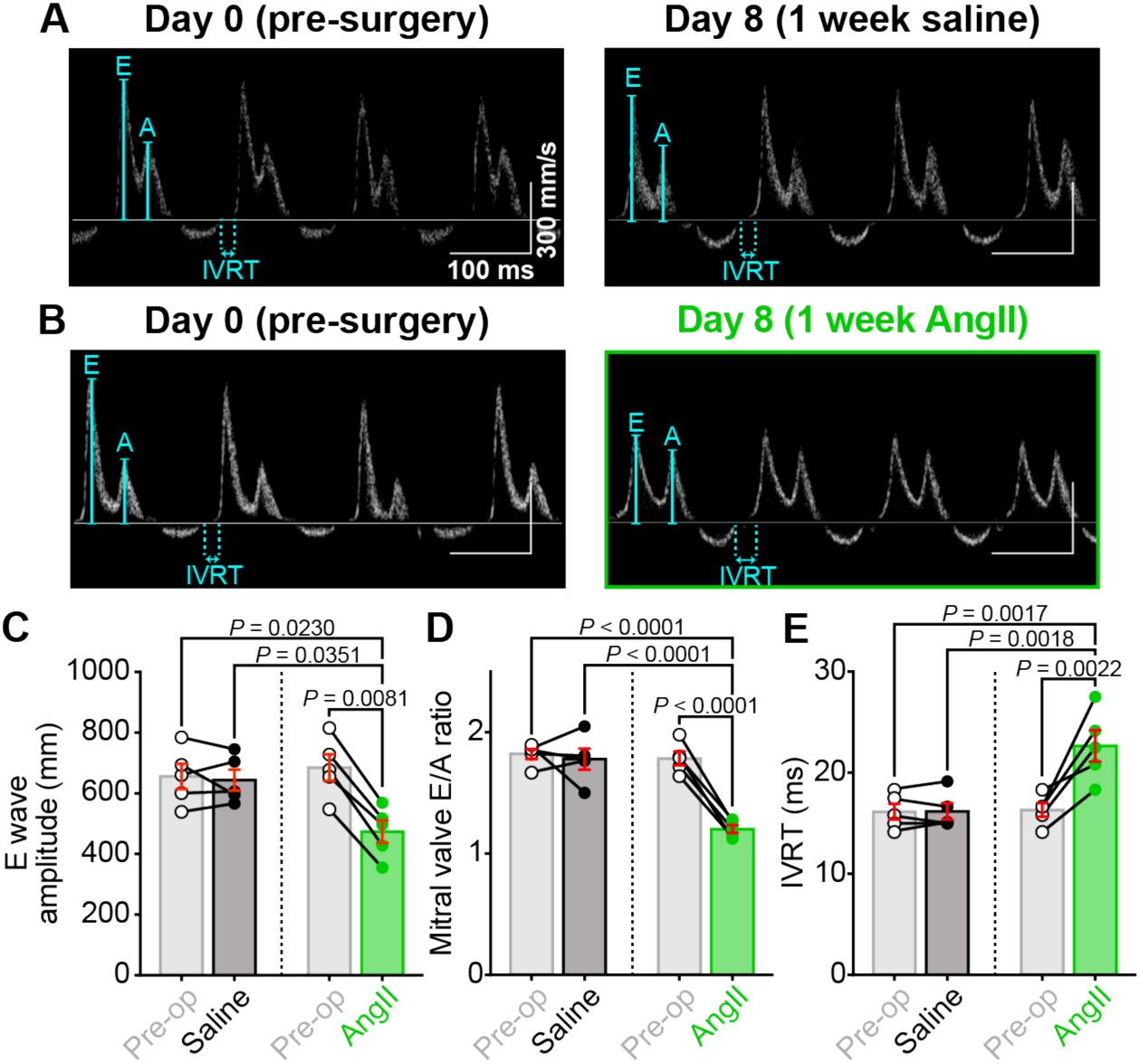
Chronic AngII infusion leads to diastolic dysfunction. Representative transmitral Doppler flow patterns from **A,** saline and **B,** AngII-infused hearts, recorded from the same mice before (pre-op, *left*) and after 1 week of infusion (post-infusion, *right*), showing early (E) and late (A) filling waves. **C-E**: Bar plots comparing paired diastolic function measurements between pre-op and post-infusion timepoints in saline (*N* = 5) and AngII-infused (*N* = 5) mice: **C,** E wave amplitude, **D,** E/A ratio, and **E,** isovolumic relaxation time (IVRT). Data analyzed using two-way ANOVA with Tukey’s post-hoc tests; error bars indicate SEM.

### T-tubule organization is disrupted by chronic AngII elevation

Chronic AngII production^8–10^ and chronic *β*-adrenergic stimulation^49, 50^ are each associated with hypertrophy and HF. The pathological mechanism that drives the transition between hypertrophy and HF is thought to involve t-tubule remodeling and disorganization^51^. Furthermore, pharmacological PIP_2_ depletion has been shown to disrupt t-tubule organization in cultured ventricular myocytes^52^. Our data suggest that chronic AngII infusion triggers PIP_2_ depletion, *β*-adrenergic stimulation, cardiac hypertrophy, and diastolic dysfunction. We thus hypothesized that chronic AngII infusion may also precipitate t-tubule disruption. To test that hypothesis, we stained live myocytes isolated from saline- and AngII-infused mice with the membrane dye, di-8-ANEPPS.

T-tubule frequency and periodicity analysis was performed using an ImageJ/Fiji plugin called TTorg^38^. This program performs a fast Fourier transform (FFT) of the confocal image of the t-tubule network. Based on the frequency of the t-tubules, a peak in the FFT spectrum develops. The amplitude of this peak is called t-tubule “power” and is used as an indication of the organizational level of the transverse-tubule network. Visually, the t-tubule staining revealed a less organized network in myocytes isolated from AngII-infused mice compared to saline controls (Figure 8A-B). Quantitative analysis confirmed this qualitative observation, revealing that t-tubule power was significantly lower in AngII-infused myocytes compared to saline controls (Figure 8C-E). Together these results reveal that t-tubule network deterioration and remodeling is stimulated by chronic AngII-infusion.

**Figure 8.**
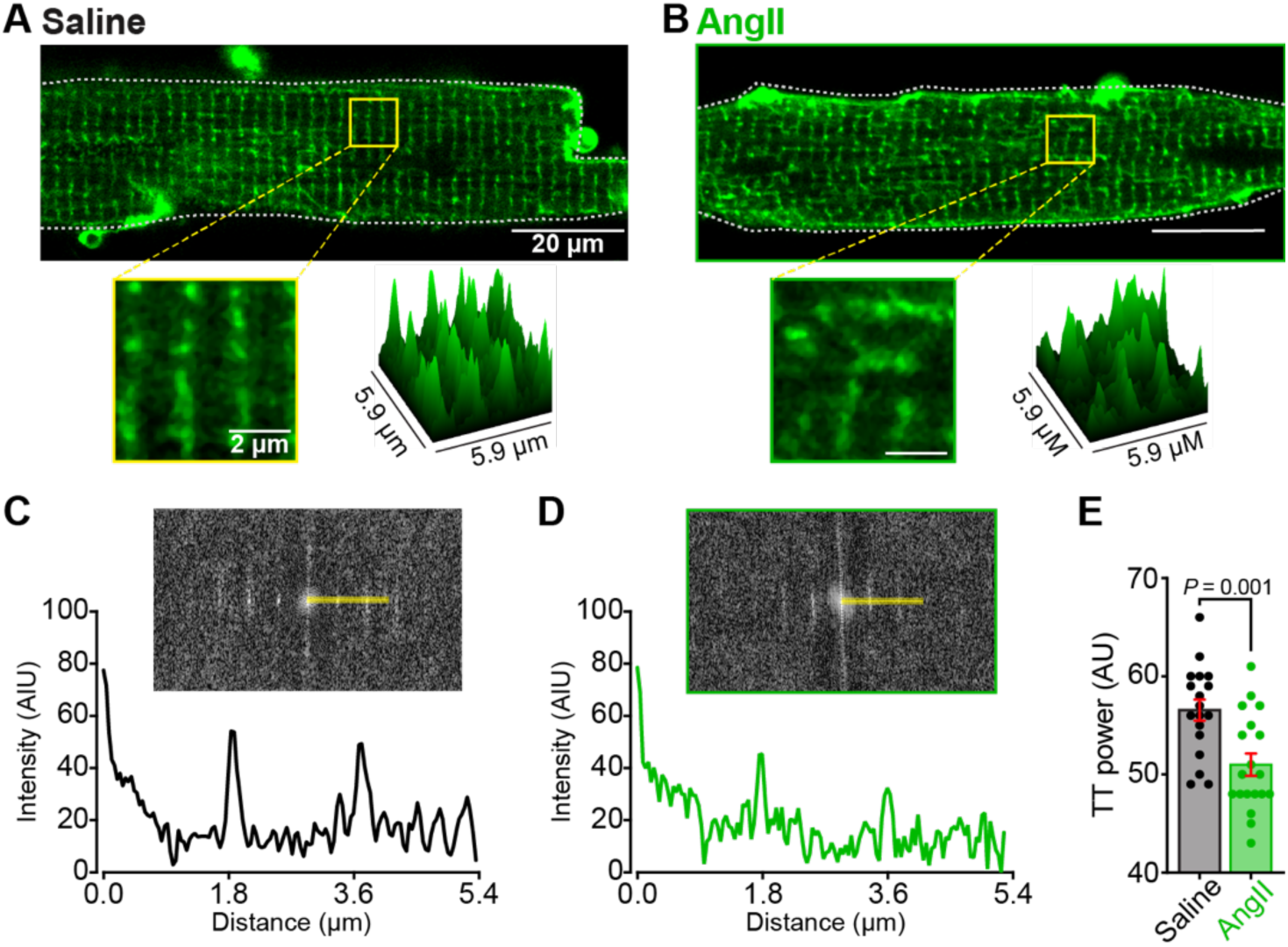
T-tubule organization is disrupted by chronic AngII stimulation. Representative Airyscan super-resolution images of myocytes from **A,** saline and **B,** AngII-infused mice stained with di-8-ANEPPS. Zoom-in and 3D surface plots of fluorescence intensity of ROIs indicated by boxes (*bottom*). Di-8-ANEPPS fluorescence intensity as a function of distance showing periodicity of t-tubule network in myocytes from **C,** saline and **D,** AngII-infused mice. **E**, Bar plot showing a t-tubule power measurements from saline (*N* = 3, *n* = 18), and AngII (*N* = 3, *n* = 19) myocytes. Statistical analysis was performed using unpaired Student’s t-test. Error bars indicate SEM.

### PI(3,4,5)P_3_ preservation does not reverse AngII-triggered alterations in I_Ca_, t-tubule organization, or cellular hypertrophy

As mentioned above, phospholipid mass spectrometry revealed that chronic AngII-infusion resulted in a 36 % reduction in PIP_3_ (phosphatidylinositol (3,4,5)-trisphosphate) in whole heart lysates. PIP_3_ is a critical player in the PI3K/Akt signaling pathway which has been shown to play several important roles in the heart including 1) enhancing Ca_V_1.2 surface expression through a mechanism involving Ca_V_β_2_ subunit phosphorylation, resulting in increased *I*_Ca_ and enhanced cardiac function^53, 54^; 2) promoting cellular hypertrophy^55^; 3) having a cardioprotective effect by inhibiting cardiomyocyte death during ischemia/reperfusion^56^ and limiting infarct size post-myocardial infarction^57^. PIP_3_ is also known to stabilize F-actin through interactions with actin-capping proteins^58^. This F-actin stabilization is crucial for maintaining t-tubule integrity in adult cardiomyocytes^59–61^. Consequently, PIP_3_ depletion leads to actin destabilization, increasing susceptibility to biomechanical stress and t-tubule loss in cardiomyocytes^58, 62, 63^. The reduced PIP_3_ levels we see with chronic AngII-infusion likely substantially diminish PI3K/Akt signaling. We thus hypothesized that this impaired signaling could explain several phenomena we observed but had yet to connect mechanistically: the decreased surface expression and clustering of Ca_V_1.2 channels, impaired cardiac function, cellular hypertrophy, and t-tubule loss in cardiomyocytes. To test these hypotheses, we utilized the PTEN inhibitor bisperoxovanadium 1,10 phenanthroline (BpV(phen))^64^ to preserve PIP_3_. PTEN is a 3’-lipid phosphatase that terminates PI3K/Akt signaling by dephosphorylating PI(3,4,5)P_3_ to form PI(4,5)P_2_, inhibiting PTEN thus elevating PIP_3_ levels and Akt signaling^65^. We infused mice with BpV(phen) (17 µg/kg/day) either on its own or in combination with AngII (1 mg/kg/day) via osmotic minipumps. Contrary to our predictions, PTEN inhibition with BpV(phen) during chronic AngII treatment failed to prevent alterations in *I*_Ca_ (Figure 9A-F), cellular hypertrophy (Figure 9G), Ca_V_1.2 channel clustering (Figure 9H), or t-tubule organization (Figure 9I-M). Since prevention of PIP_3_ breakdown did not attenuate or negate the chronic AngII-triggered alterations in *I*_Ca_, Ca_V_1.2 clustering and expression, t-tubule organization, or cellular hypertrophy, these results reveal that reduced PIP_3_ is not responsible for these aberrant changes.

**Figure 9.**
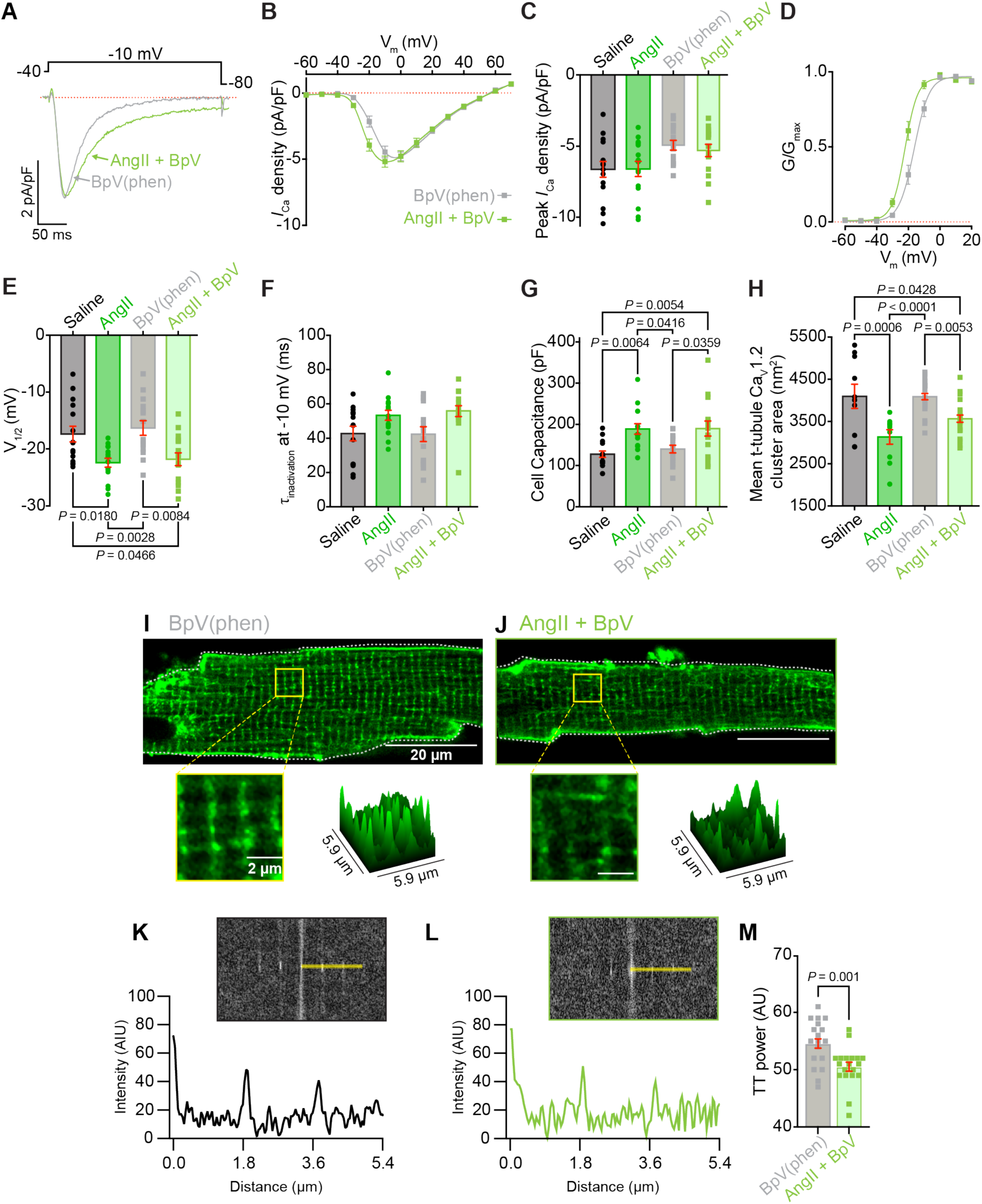
PTEN inhibition does not prevent AngII-induced changes in *I*_Ca_ or cellular architecture. **A,** Representative whole-cell *I*_Ca_ during 300 ms depolarization to −10 mV, and **B,** corresponding current-voltage relationships from myocytes isolated from BpV(phen)-infused (grey) and AngII+BpV(phen)-infused (light-green) mice. **C-H,** Comparison of electrophysiological and structural parameters between BpV(phen) (*N* = 4, *n* = 15) and AngII+BpV(phen) (*N* = 4, *n* = 15) groups, shown alongside saline and AngII-infused data from Figure 4 for comparison when necessary: **C,** peak *I*_Ca_ density; **D,** voltage-dependence of normalized conductance (G/G_max_) with Boltzmann fits; **E,** half-maximal activation voltage (V_1/2_); **F,** time constant of *I*_Ca_ inactivation at −10 mV; and **G,** cell capacitance. **H,** Ca_V_1.2 cluster area from SMLM analysis of BpV(phen) (*N* = 4, *n* = 24) and AngII+BpV(phen) (*N* = 4, *n* = 23) groups, shown alongside saline and AngII-infused data from Figure 2. **I-J,** Representative di-8-ANEPPS staining of t-tubules in **I,** BpV(phen)-infused and **J,** AngII+BpV(phen)-infused myocytes. **K-M,** T-tubule organization analysis showing di-8-ANEPPS fluorescence intensity versus distance in **K,** BpV(phen) and **L,** AngII+BpV(phen)-infused myocytes, with **M,** bar plot summarizing t-tubule power measurements from BpV(phen) (*N* = 4, *n* = 21) and AngII+BpV(phen) (*N* = 4, *n* = 19) groups. Data were analyzed using two-way ANOVA with Tukey’s post-hoc tests (**C, E-H**) or unpaired Student’s t-test (**M**). Error bars indicate SEM.

### PI(3,4,5)P_3_ preservation prevents the damaging effects of chronic AngII on cardiac function

We also examined *in vivo* cardiac function in mice performing echocardiography and doppler imaging before and after one-week infusion with BpV(phen) or BpV(phen) plus AngII. Despite the maintained cardiomyocyte and *I*_Ca_ remodeling observed in isolated ventricular myocytes, we observed that BpV(phen) either blunted or completely abolished the negative effects of AngII on systolic and diastolic function when the two were co-infused (Figure 10, Figure 11, and Figure S1). The cardiac hypertrophy that was evident in the measurements made from AngII-infused animals did not manifest when the PTEN-inhibitor was co-infused. Given that the co-infusion did not correct the myocyte remodeling, these results suggest that PTEN inhibition may prevent cardiac fibrosis and could provide a level of cardioprotection in hypertensive patients.

**Figure 10.**
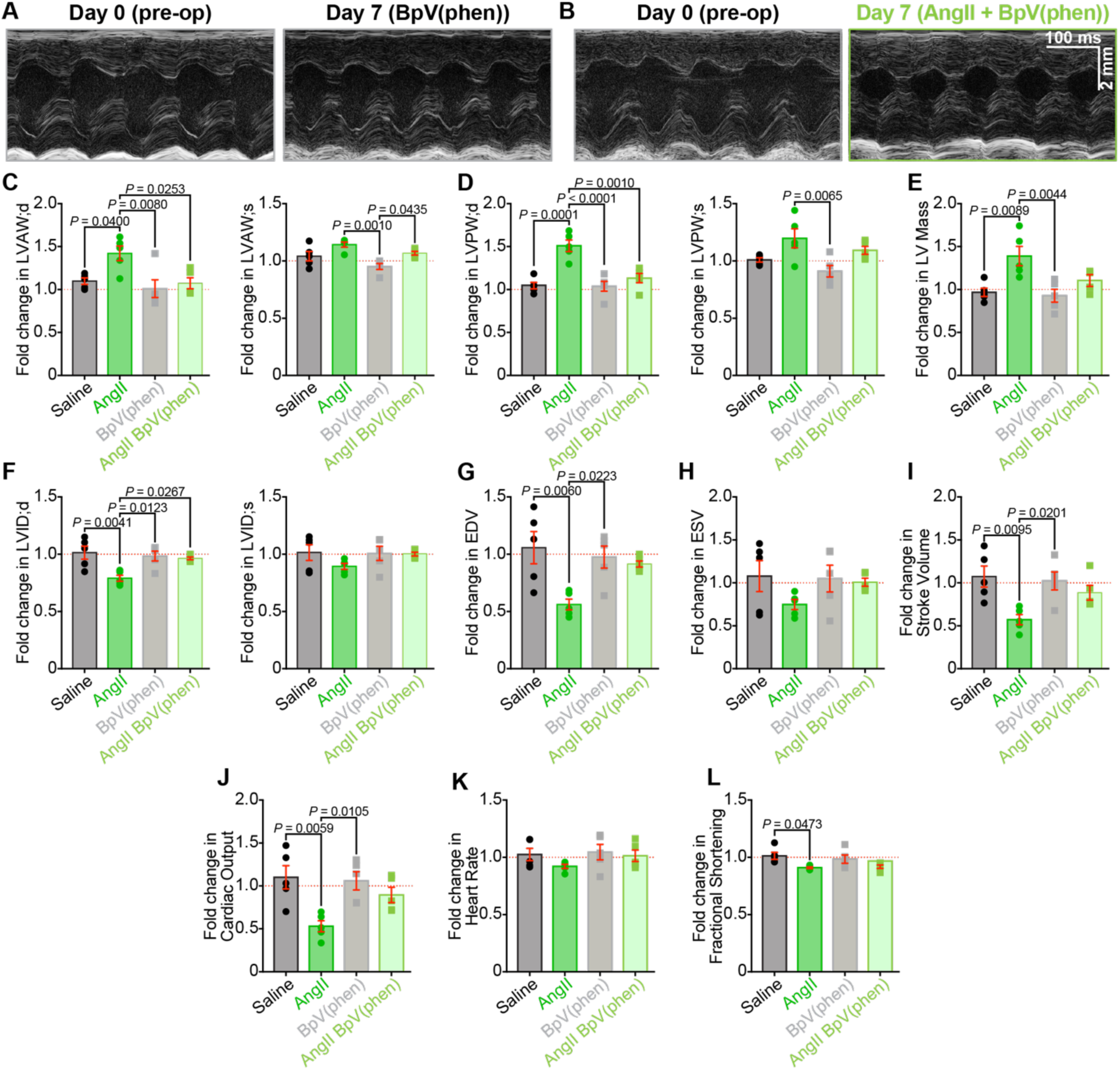
PTEN inhibition prevents AngII-induced structural remodeling. Representative short-axis M-mode echocardiograms from **A,** BpV(phen) and **B,** BpV(phen)+AngII-treated hearts, recorded from the same mice before (pre-op, *left*) and after 1 week of infusion (post-infusion, *right*). **C-L,** Bar plots comparing paired echocardiographic parameters between pre-op and post-infusion timepoints in BpV(phen) and BpV(phen)+AngII-treated mice (*N* = 5/group), shown alongside the saline and AngII-infused data from Figure 6 for comparison: **C,** left ventricular anterior wall thickness and **D,** posterior wall thickness in diastole (*left*) and systole (*right*); **E,** left ventricular mass (LV mass); **F,** left ventricular inner diameter (LVID); **G,** end diastolic volume (EDV); **H,** end systolic volume (ESV); **I,** stroke volume; **J,** cardiac output; **K,** heart rate; and **L,** fractional shortening. Data were analyzed using two-way ANOVAs with Tukey’s multiple comparisons post hoc test. Error bars indicate SEM.

**Figure 11.**
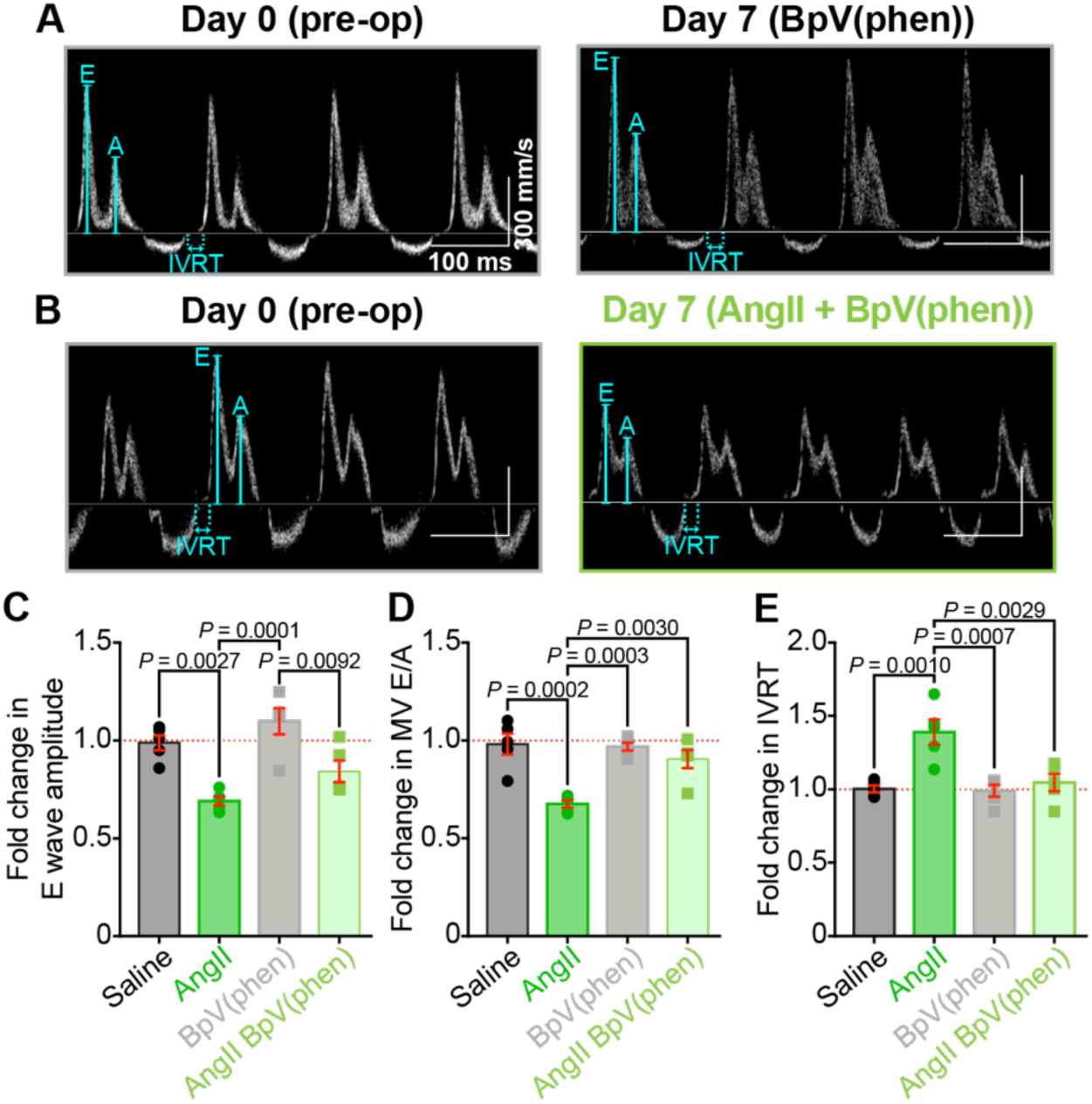
PTEN inhibition preserves diastolic function during AngII treatment. Representative transmitral Doppler flow patterns from **A,** BpV(phen) and **B,** BpV(phen)+AngII-treated hearts, recorded from the same mice before (pre-op, *left*) and after 1 week of infusion (post-infusion, *right*). **C-E,** Bar plots comparing paired diastolic function measurements between pre-op and post-infusion timepoints in BpV(phen) and BpV(phen)+AngII-treated mice (*N* = 5/group), shown alongside the saline and AngII-infused data from Figure 7 for comparison: **C,** E wave amplitude, **D,** E/A ratio, and **E,** isovolumic relaxation time (IVRT). Data were analyzed using two-way ANOVAs with Tukey’s multiple comparisons post hoc test. Error bars indicate SEM.

## Discussion

Our investigation into chronic AngII signaling reveals a complex interplay between cardiac phosphoinositides, ion channel regulation, and pathological remodeling. We demonstrate that sustained AngII exposure profoundly alters the cardiac phosphoinositide landscape, with widespread implications for the many proteins regulated by these lipids. Building on our previous work showing acute AngII-induced PI(4,5)P_2_ depletion triggers Ca_V_1.2 channel internalization to early endosomes, we now show that chronic AngII exposure results in broader redistribution of channels throughout the endosomal network, including early, recycling, and late endosomal compartments. Although this endocytosis reduces t-tubular Ca_V_1.2 clustering and expression, enhanced sympathetic drive maintains Ca^2+^ signaling by increasing channel phosphorylation. However, this compensation proves insufficient to prevent cardiac dysfunction, which develops alongside pathological hypertrophy and fibrosis. Importantly, we identify PIP_3_ depletion as a critical mediator of this pathological remodeling. Preventing PIP_3_ loss through PTEN inhibition reduces left ventricular hypertrophy and preserves cardiac function without affecting myocyte Ca^2+^ handling or cytoarchitecture, suggesting its effects occur primarily through reduction of fibrosis. These findings establish PIP_3_ maintenance as a promising therapeutic strategy for hypertensive heart disease and as a potential protective adjunct therapy during clinical AngII administration.

Our finding that the t-tubular Ca_V_1.2 population is selectively vulnerable to AngII-induced internalization, while surface/crest populations remain intact, aligns with previous observations by Hermosilla et al., who reported similar compartment-specific channel removal following 30-60 minute AngII exposure in isolated cardiomyocytes^66^. This spatial specificity suggests differential AT_1_R expression or distinct distribution of PLC effector enzyme between t-tubular and crest membranes, although direct confirmation awaits development of validated AT_1_R antibodies. Furthermore, we found that chronic AngII infusion negatively affects the t-tubule network and cytoarchitecture; thus, it is also possible that detubulation is at least a partial factor in the loss of the t-tubule population of Ca_V_1.2. However, multiple lines of evidence support direct channel internalization as the primary mechanism. Prior studies in both cardiomyocytes and HEK-293 cells demonstrated that AT_1_R stimulation triggers channel endocytosis^66^ and accumulation in early endosomes^29^. Extending these findings, our chronic AngII infusion model reveals even more extensive channel removal from the sarcolemma and accumulation throughout the endosomal network, including early, recycling, and late endosomal compartments. Some of these channels may ultimately be directed toward degradation.

Unlike studies of ischemic and dilated cardiomyopathy, where t-tubule remodeling leads to redistribution of Ca_V_1.2 channels from t-tubules to surface crests^12, 67, 68^, we observed no compensatory increase in crest channel density. This difference may reflect distinct temporal phases of cardiac remodeling, with our one-week AngII infusion model capturing an earlier stage before channel redistribution occurs. Despite reduced t-tubular Ca_V_1.2 expression, whole-cell current amplitude remained comparable to saline-infused controls, suggesting functional upregulation of the remaining channels.

This upregulation likely occurs through enhanced sympathetic signaling, as chronic AngII can increase sympathetic outflow via activation of AT_1_Rs in the central nervous system^69–71^. Chronic AngII signaling can also facilitate sympathetic neurotransmission by stimulating release and inhibiting reuptake of norepinephrine^72–74^, which in turn can activate PKA and CaMKII downstream of *β*-AR activation. Indeed, heart lysates from AngII-infused mice showed increased phosphorylation of PKA targets (Ca_V_1.2-Ser1928, RyR2-Ser2808) and the CaMKII target RyR2-Ser2814, indicating activation of both kinase pathways. While previous studies have linked CaMKII-phosphorylation to enhanced crest Ca_V_1.2 activity in dilated cardiomyopathy and PKA-phosphorylation to t-tubular Ca_V_1.2 in ischemic cardiomyopathy^67^, our study suggests both kinases may contribute to channel regulation during chronic AngII signaling.

Interestingly, *I*_Ca_ recorded from ventricular myocytes isolated from AngII-infused mice showed slower inactivation kinetics compared to controls. Though *β*-adrenergic stimulation typically reduces voltage-dependent inactivation (VDI)^39–41, 43^, it usually accelerates overall *I*_Ca_ inactivation through increased Ca^2+^-dependent inactivation (CDI) due to enhanced channel open probability and calcium-induced calcium release (CICR)^43^. This raises a crucial question: why doesn’t chronic AngII infusion, which produces sympathetic stimulation, accelerate *I*_Ca_ inactivation? We propose that the answer lies in the channel localization since channel location also influences inactivation kinetics. T-tubule-localized channels within dyadic cleft microdomains experience rapid CDI due to confined Ca^2+^ accumulation, whereas crest-localized channels exhibit slower inactivation due to less restricted Ca^2+^ diffusion^67^. Thus, the preserved *I*_Ca_ magnitude but slowed inactivation we observed with chronic AngII likely reflects both altered channel distribution and phosphorylation state. The preferential loss of rapidly inactivating t-tubule Ca_V_1.2 channels leaves a greater relative contribution from slowly inactivating crest channels, which could prolong action potential duration and ultimately become arrhythmogenic.

The t-tubule localized sub-population of Ca_V_1.2 channels are most important for excitation-contraction coupling. Despite reduced t-tubular channel density, Ca^2+^ transient magnitude was maintained and slightly enhanced during chronic AngII infusion, suggesting that phosphorylation of both remaining Ca_V_1.2 channels and RyR2s effectively preserved EC-coupling, even though echocardiography revealed impaired cardiac function *in vivo*. While this compensatory phosphorylation successfully maintained EC-coupling in our one-week AngII model, sustained RyR2 hyperphosphorylation in sustained chronic hypertension can become progressively maladaptive. The resulting increase in RyR2 leak elevates diastolic [Ca^2+^]_i_ and triggers hypertrophic signaling^75^, suggesting our findings may capture an early adaptive phase. The end stage of hypertensive heart disease is dilated cardiomyopathy^76^ with diastolic dysfunction and reduced ejection fraction. Our echocardiography reveals that one week of chronic AngII infusion had not progressed to end-stage but is more akin to a degree II stage of hypertensive heart disease with LV diastolic dysfunction and LV hypertrophy.

Intriguingly, while *β*-adrenergic signaling promotes Ca_V_1.2 channel recycling to t-tubule membranes^77, 78^, we found that chronic AngII infusion drove channel endocytosis despite concurrent *β*-adrenergic activation, suggesting AngII’s effects on trafficking override *β*-adrenergic recycling signals. This trafficking imbalance may stem from the widespread alterations in cardiac phosphoinositide (PI) levels we observed, as these lipids orchestrate endosomal trafficking^79^. Each step of membrane protein trafficking requires specific PI species: PI(4,5)P_2_ clustering and conversion to PI(4)P enables endocytosis^80^, while endosomal PI(3)P recruits sorting proteins that direct cargo fate^81^. Indeed, disrupting PI(3)P homeostasis through MTM1 knockdown has been found to reduce cargo recycling^82^, highlighting the importance of precise PI regulation. While our mass spectrometry detected alterations in many PI species with chronic AngII, including PIP, further studies are needed to determine how specific PIs contribute to altered Ca_V_1.2 trafficking and cardiac remodeling.

The significant reduction in cardiac PIP_3_ levels following chronic AngII infusion drew our attention given PIP_3_’s central role in PI3K/Akt signaling and its regulatory effects on Ca_V_1.2 channels. Class I phosphoinositide 3-kinases (PI3K) generate PIP_3_ by phosphorylating plasma membrane PI(4,5)P_2_ at the 3’ position of the inositol ring. PIP_3_ then recruits Akt (protein kinase B) to the membrane through its pleckstrin-homology (PH) domain, where Akt is activated by mTORC2 and PDK1-mediated phosphorylation^83^. Activated Akt modulates multiple downstream pathways, including promotion of Ca_V_1.2 channel surface expression in neurons^84^, smooth muscle cells^85^, and cardiomyocytes^53, 83^ through Ca_V_β subunit phosphorylation, ultimately increasing *I*_Ca_. This signaling cascade is terminated by PTEN (phosphatase and tensin homolog), which dephosphorylates PIP_3_ to PIP_2_. The importance of this pathway for cardiac function has been demonstrated in genetic models where cardiac-specific PTEN deletion increases Akt activity and *I*_Ca_ ^54^, while PI3Kα-null myocytes show reduced *I*_Ca_ that can be rescued by PIP_3_ supplementation to improve cardiac performance^86^.

PI3K/Akt signaling is also well known to affect cell growth and hypertrophy. Mammalian target of rapamycin (mTOR) is a downstream target of Akt and mTOR inhibition with rapamycin can prevent cardiac hypertrophy^87^. In our hands, PTEN inhibition neither prevented AngII-induced cellular hypertrophy nor protected Ca_V_1.2 channel expression and *I*_Ca_. Instead, it provided broad cardioprotection during AngII infusion through apparent non-myocyte mechanisms. This protection manifested as preserved cardiac structure (LV mass and dimensions) and function (stroke volume, cardiac output, and fractional shortening), along with improved diastolic function (preserved E/A ratio and IVRT) compared to AngII treatment alone. The disconnect between persistent cellular hypertrophy and improved cardiac function with PTEN inhibition suggests effects primarily on the cardiac interstitium rather than cardiomyocytes. Indeed, previous studies have shown that PTEN inhibition provides “fibro-protection” by promoting anti-inflammatory M2 macrophage activation in both spontaneously hypertensive rats and AngII-induced hypertension models^88^. This anti-fibrotic effect, observed with either PTEN inhibition or Akt siRNA, appears to be mediated through a PI3K/Akt/TGF-β/Smad-2/3 pathway.

The profound changes in PI homeostasis caused by chronic AngII infusion may reflect changes in cellular energetics. Most lipid kinases require ATP as a phosphate donor for their catalytic function, suggesting that changes in cellular ATP availability could explain the widespread alterations we observe in various PI species. Recent findings have revealed that bulk cytosolic ATP concentrations are considerably lower than previously estimated, with diastolic levels measured at ∼450 µM^89^. This has important implications for PI homeostasis during heart failure, where impaired energy metabolism can reduce cardiac ATP content by up to 30% compared to healthy hearts^90^. While the K_m, ATP_ of most lipid kinases lies in the 5-100 µM range^91^, local ATP depletion during pathological conditions could decrease ATP levels below the affinity range of key lipid kinases, compromising their catalytic activity. Such ATP deficits could be particularly relevant for PI3K function during heart failure, potentially compounding the PIP_3_ depletion we observe with chronic AngII stimulation. This represents an unappreciated mechanism linking metabolic dysfunction to impaired PI3K/Akt signaling in cardiac pathology. The relationship between cellular energetics and phosphoinositide signaling suggests that therapeutic strategies to preserve both ATP levels and PIP_3_ (through PTEN inhibition) may provide enhanced cardioprotection by maintaining PI homeostasis during disease progression.

Our study reveals how chronic AngII signaling orchestrates a complex series of molecular events in the heart, from phosphoinositide remodeling to ion channel trafficking and ultimately cardiac dysfunction. We demonstrate that while enhanced sympathetic drive can temporarily compensate for reduced t-tubular Ca_V_1.2 expression through channel phosphorylation, this adaptation proves insufficient to prevent pathological remodeling. Most significantly, we identify PIP_3_ depletion as a critical mediator of AngII-induced cardiac dysfunction and demonstrate that preventing PIP_3_ loss through PTEN inhibition provides substantial cardioprotection. The fact that PTEN inhibition preserves cardiac function without affecting cellular hypertrophy or Ca_V_1.2 trafficking suggests it acts primarily through non-myocyte mechanisms, likely by reducing fibrosis via PI3K/Akt/TGF-β signaling in cardiac fibroblasts. These findings not only advance our understanding of hypertensive heart disease progression but also identify PTEN inhibition as a promising therapeutic strategy. The cardioprotective effects of PTEN inhibition during chronic AngII exposure suggest it could potentially serve as an adjunct therapy in conditions requiring clinical AngII administration or in patients with hypertension-induced cardiac remodeling.

## Acknowledgements

We are grateful to Mr. Joshua Tulman who produced the artwork featured in the Graphical Abstract.

## Non-standard Abbreviations and Acronyms

AngII: Angiotensin II
AP: Action potential
AT1R: Angiotensin II type 1 receptor
AT2R: Angiotensin II type 2 receptor
BpV(phen): Bisperoxovanadium 1,10 phenanthroline
CaMKII: Ca2+/calmodulin-dependent protein kinase II
CDI: Calcium-dependent inactivation
EC: Excitation-contraction
EDV: End diastolic volume
ESV: End systolic volume
GLOX: Glucose oxidase
HF: Heart failure
HW/BW: Heart weight to body weight ratio
ICa: Calcium current
IVRT: Isovolumetric relaxation time
LA: Left atrium
LV: Left ventricle/left ventricular
LVAW: Left ventricular anterior wall
LVID: Left ventricular inner diameter
LVPW: Left ventricular posterior wall
MEA: Cysteamine
mTOR: Mammalian target of rapamycin
PI: Phosphoinositide
PIP_2_: Phosphatidylinositol 4,5-bisphosphate
PIP_3_: Phosphatidylinositol 3,4,5-trisphosphate
PKA: Protein kinase A
PLC_ε_: Phospholipase C epsilon
PTEN: Phosphatase and tensin homolog
RAAS: Renin-Angiotensin-Aldosterone-System
SMLM: Single-molecule localization microscopy
SR: Sarcoplasmic reticulum
VDI: Voltage-dependent inactivation

